# Voluntary engagement predicts recovery across rehabilitation time points after experimental spinal cord injury

**DOI:** 10.64898/2026.09.28.751615

**Authors:** Jarred M. Griffin, Panagiotis Papaioannou, Rohan Menon, Olivia Cheyne, Simon Kellaway, Darren Svirskis, Catherine Jutzeler, Ahmad Aziz, Frank Bradke

## Abstract

Rehabilitation is the cornerstone of clinical care following spinal cord injury (SCI) and is frequently incorporated into preclinical studies to evaluate emerging therapies. However, the variables that influence the effectiveness of rehabilitation interventions in rodent models remain poorly characterized, limiting the translational value of preclinical research. Here, we performed a meta-analysis of 150 preclinical studies comprising more than 3000 rodents and 292 effect sizes to identify such factors associated with successful rehabilitation after SCI. Rehabilitation improved functional outcomes across multiple behavioral domains, with larger effects observed in thoracic than cervical injury models. Meta-regression identified environmental enrichment as the most effective strategy, particularly when combined with task-specific training. Longer rehabilitation duration was associated with improved recovery, whereas greater daily exposure to forced rehabilitation was associated with smaller gains. In contrast, prolonged access to voluntary rehabilitation did not show this relationship. Rehabilitation improved outcomes when initiated at all time points after injury, with greater benefits when initiated during the late subacute and intermediate phases. Rehabilitation approaches associated with higher stress burden produced reduced benefits, suggesting that stress may limit rehabilitation-induced recovery. Collectively, these findings identify rehabilitation type, duration, and delivery context as important determinants of recovery after SCI and support a broad therapeutic window for intervention. Our results favor low-stress, voluntary rehabilitation strategies that promote sustained engagement and provide principles for designing effective and biologically appropriate rehabilitation strategies, potentially improving translational success for emerging SCI therapies.

**One Sentence Summary:** Sustained voluntary rehabilitation promotes functional recovery across stages of experimental spinal cord injury, whereas prolonged forced training may reduce its benefits.

## INTRODUCTION

Despite advances in basic and preclinical research that provide promising therapeutic perspectives, spinal cord injury (SCI) still lacks an approved regenerative treatment (*1-9*). Beyond early decompressive surgery, rehabilitation remains the only intervention that consistently improves function in patients (*10*). Clinical rehabilitation encompasses diverse goals, including independence, compensation, and quality of life, whereas preclinical rehabilitation is primarily designed with the intention to enhance functional recovery and activity-dependent neuroplasticity (*11, 12*). Recognizing these distinctions is important when evaluating regenerative therapies in combination with rehabilitative interventions.

Animal models have played a critical role in reproducing rehabilitation-induced recovery observed in patients, enabling investigation of the mechanisms underlying functional improvements and assessment of rehabilitation as part of combinatorial therapeutic strategies (*6*). Over the years, a wide range of rehabilitation paradigms have been developed in animal studies, varying in training type, task specificity, onset, duration, and intensity (*6, 11, 12*). Likewise, functional recovery is assessed using a diverse array of behavioral outcome measures (*13*).

While this heterogeneity complicates direct comparisons between studies, it also provides an opportunity to identify the rehabilitation parameters most strongly associated with recovery through quantitative synthesis of the available evidence. In fact, meta-analyses of published studies have helped to discover therapeutic treatments and intervention strategies in the areas of multiple sclerosis (*14, 15*), cognitive impairment (*16*) and oncology (*17*).

For SCI, previous systematic reviews and meta-analyses have concluded that rehabilitative interventions can promote functional recovery (*18-23*). However, the key factors driving rehabilitation-induced improvements remain poorly understood in both clinical and preclinical settings (*24, 25*). Thus, critical aspects of rehabilitation delivery, including optimal timing, duration, and overall training dose, remain largely undefined (*26*).

To address these knowledge gaps, we performed a comprehensive meta-analysis and meta-regression of experimental SCI studies to determine how rehabilitation type, initiation timing, duration, and delivery characteristics influence functional recovery. Our results uncover the key drivers of rehabilitation-induced recovery in rodent models. These findings provide a strategy for aligning regenerative treatments with optimal rehabilitation paradigms, thereby improving the translational potential of preclinical research for patients.

## RESULTS

### Study design and characteristics of the analyzed dataset

We started by generating a list of studies from the PubMed database with the search term “spinal cord injury” and “rehabilitation” and “rat” or “mouse” (specific search terms are included in Methods) for the publication interval 2000-2025. The search resulted in 441 articles that were subject to inclusion screening (**Fig. 1A**). Studies were included if they reported behavioral data, defined SCI groups, enabled data extraction from the figures, tables or other supplementary resources, and include control groups (i.e., rehabilitation without additional intervention). A total of 150 published studies, reporting 292 effect size calculations, met the eligibility criteria and were included in the meta-analysis (**Fig. 1A**). Collectively, this encompassed data from over 3000 adult mice and rats. Detailed characteristics, demographics, and citations of the included studies are provided in the supplementary data file (**Data File S1**).

**Fig. 1.**
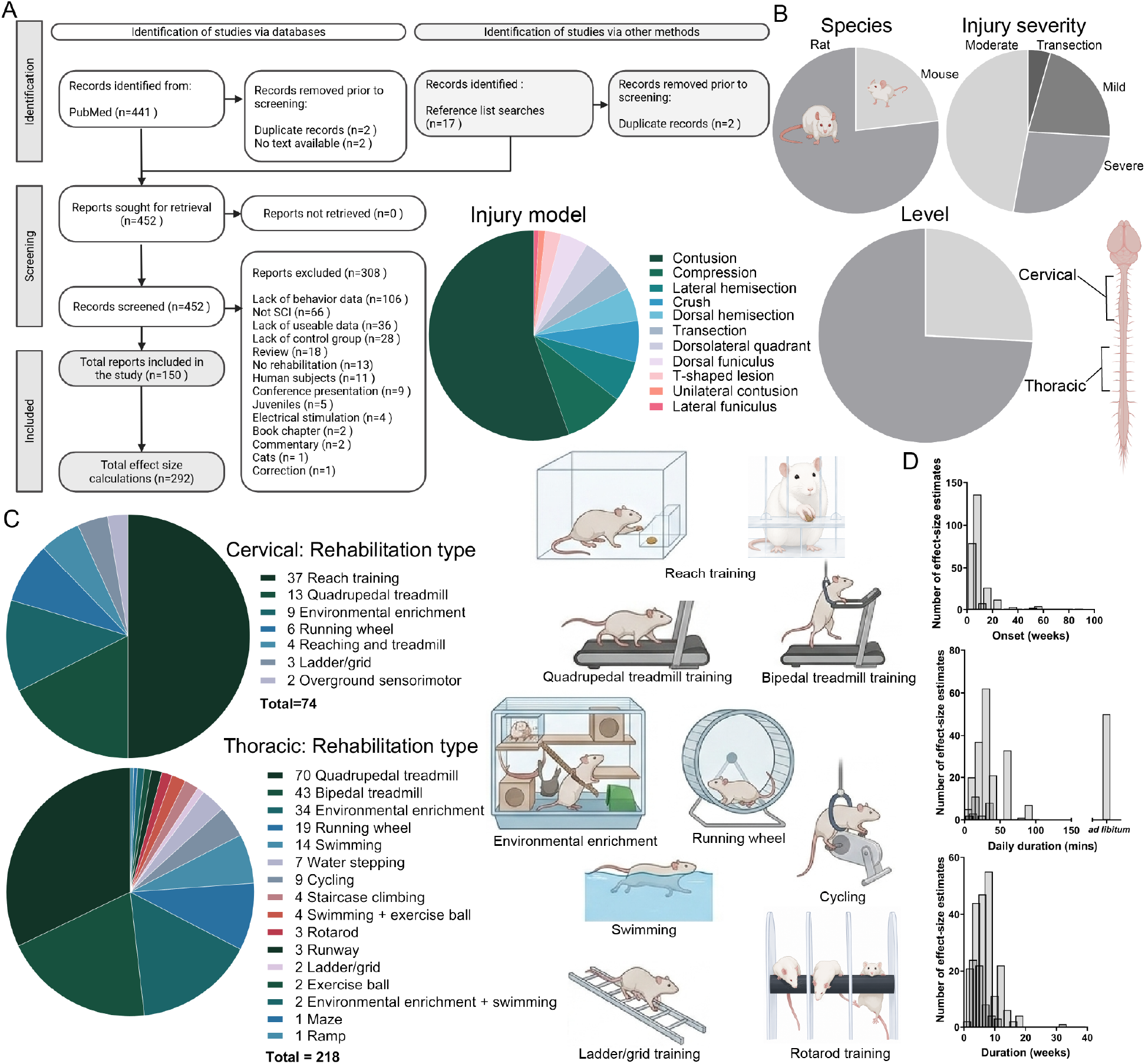
Study selection, experimental characteristics, and rehabilitation parameters included in the meta-analysis. **(A)** PRISMA flow diagram summarizing study identification, screening, eligibility, and inclusion. A total of 441 records were identified through database searches and 17 through reference lists; after removal of duplicates and exclusions, 150 studies were included, yielding 292 effect size calculations. **(B)** Distribution of included studies by species (rat vs. mouse), injury severity (moderate, severe, mild, transection), injury model, and injury level (thoracic vs cervical). **(C)** Distribution of rehabilitation modalities used in cervical and thoracic injury models. Reach training was the predominant intervention in cervical models, whereas quadrupedal and bipedal treadmill training were most common in thoracic models. Includes representative illustrations of common rehabilitation paradigms used across studies, including reach training, running wheel, treadmill training (quadrupedal and bipedal), cycling, environmental enrichment, ladder training, swimming, and rotarod training, as well as complex multifactorial enrichment systems. **(D)** Histograms of daily onset timing (weeks post-injury), training duration (minutes), and overall intervention duration (weeks), with separate indication of *ad libitum* (continuous voluntary) exposure.

The studies represent a range of SCI models and injury levels, as well as variations in severity (**Fig. 1B**). In the studies included, 77% used rats as the experimental model, while 23% used mice (**Fig. 1B**). The severity of the injuries was categorized (**see Methods**) into complete transection (4%), severe (27%), moderate (47%) or mild (22%). 74% of studies reported thoracic-level injuries, whereas the remaining 26% used cervical-level injuries. The majority of these injury models (**Fig. 1B**) were contusions (55%), followed by compression injury (9%), lateral hemisection, crush (each 6%), and dorsal hemisection (5%).

In total, 19 different rehabilitation strategies were identified, and these were separated by the level of injury used in the experimental model. For cervical-level injuries, 50% of the studies used reach training as the rehabilitation type (**Fig. 1C**). This was followed by quadrupedal treadmill training (18%), environmental enrichment (12%), and running wheel (8%). For thoracic-level injuries, the most common methods were quadrupedal treadmill training (32%), bipedal treadmill training (20%), environmental enrichment-based rehabilitation (16%), followed by running wheel (9%) (**Fig. 1C**). We further subclassified rehabilitation types based on the following treadmill training criteria: utilization of weight support, forced or voluntary training, and theincorporation of task-specific elements into environmental enrichment to distinguish between ‘targeted environmental enrichment’ and ‘general environmental enrichment’ (**Data File S1**).

For most studies (91%), rehabilitation intervention began within three weeks post-injury, with 50% initiating rehabilitation seven days after injury (**Fig. 1D**). The common daily rehabilitation duration for most regimes was 25 minutes. However, for studies where the rehabilitation was included within their home cage as in the majority of environmental enrichment or voluntary wheel running, for example, daily rehabilitation was then represented separately as ‘*ad libitum’* (**Fig. 1D**). The overall rehabilitation period across studies was between approximately four to six weeks (41%), with some protocols extending up to 32 weeks (0.4%) (**Fig. 1D**). This curated data set was the basis to uncover the key parameters that determine the effectiveness of rehabilitation.

### Rehabilitation differentially impacts recovery following cervical and thoracic spinal cord injury

Using our curated data set (**Data File S1**), we first asked whether the inclusion of a rehabilitative intervention leads to improvements in behavioral outcomes. Given the behavioral and regenerative differences between cervical and thoracic SCIs (*27, 28*), we analyzed these as separate datasets. To enable comparisons, we restricted our analysis to behavioral measures that are relatively standardized across laboratories (**Table S1**) and only included outcomes reported in at least two independent studies.

We report standardized mean differences (SMDs) together with their 95% confidence intervals (CIs) to express differences between control and rehabilitation groups in standard deviation units. Because individual studies often contributed multiple effect-size estimates, these estimates could not be considered statistically independent. We therefore used a multi-level, random-effects meta-analysis to account for their dependency (see **Methods, Fig. 2**). By modelling effect sizes as nested within studies, this approach preserves the multilevel structure of the data and enables valid comparisons between experimental groups. It also reduces the risk of inflated Type I error rates that can arise when dependent effect sizes are incorrectly treated as independent (*29*).

**Fig. 2.**
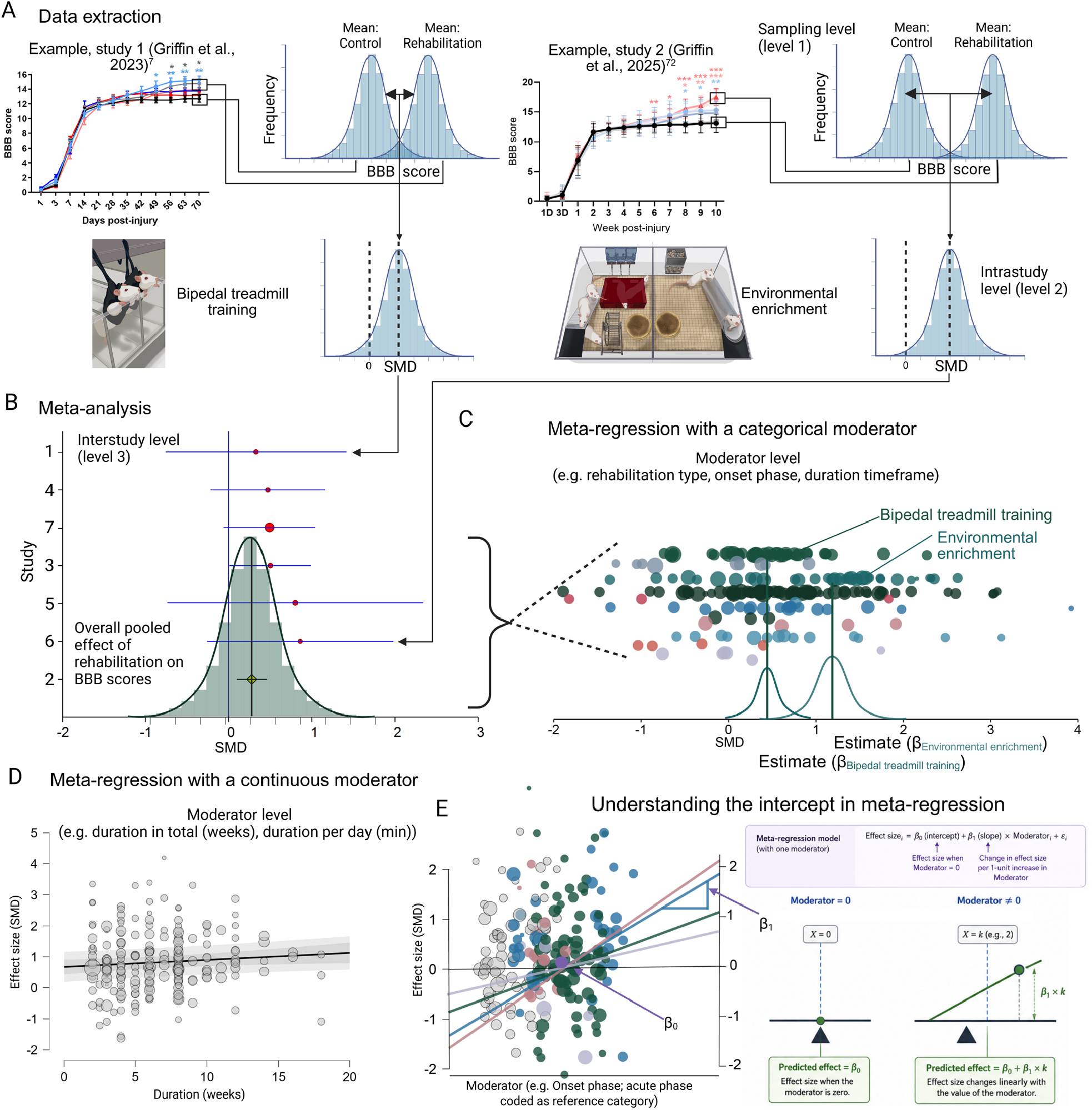
Overview of the meta-analytic workflow and meta-regression approach used to evaluate rehabilitation efficacy following spinal cord injury. **(A)** Data extraction pipeline. Behavioral outcomes were extracted at the final assessment timepoint from individual studies (examples given are: (*7, 72*)) and converted to SMDs between rehabilitation and control groups. Multiple effect sizes could be derived from a single study based on rehabilitation modality and outcome measure, creating a hierarchical data structure consisting of sampling-level, within-study, and between-study variance components. **(B)** Random-effects meta-analysis. Individual study effect sizes were pooled to generate an overall estimate of rehabilitation efficacy while accounting for heterogeneity across studies. Forest plot and density distribution illustrate the calculation of the pooled effect size and corresponding uncertainty. **(C)** Meta-regression using categorical moderators. Categorical variables, including rehabilitation type, rehabilitation onset phase, and rehabilitation-duration categories, were used to determine whether treatment efficacy differed among predefined groups. Effect estimates are generated relative to either individual subgroup means (no-intercept model) or a specified reference category (intercept model). **(D)** Meta-regression using continuous moderators. Continuous variables, including total rehabilitation duration (weeks) and daily rehabilitation exposure (minutes/day), were evaluated to determine whether treatment efficacy changed as a function of rehabilitation dose. **(E)** Conceptual representation of meta-regression. The slope coefficient (β_1_) quantifies the change in treatment effect associated with variation in the moderator, whereas the intercept (β_0_) represents the predicted effect size when the moderator equals zero or, for categorical variables, the effect size of the reference category. **(F)** Interpretation of intercept-based meta-regression models. In categorical meta-regression, the intercept represents the estimated effect size of the reference group, while subsequent coefficients represent differences between comparison groups and the reference category. Positive and negative coefficients indicate increases or decreases in treatment efficacy relative to the reference group, respectively. Created using data from the present systematic review and meta-analysis.

For initial data examination we used descriptive statistics, quantile-quantile (Q-Q) plots, meta-analytical readouts and funnel plots (**Fig. S1**). Outliers were identified with standardized effect size (*z*_*i*_) scores ± 2.5. These analyses indicated three outliers: two in the thoracic dataset (*30, 31*) and one in the cervical dataset (*32*) (**Figs. S1C-E**). Hence, heterogeneity was high in both datasets prior to outlier removal (thoracic: *Q*_*E*_ (217) = 611.72, *P* <.001; cervical: *Q*_*E*_ (73) = 134.00, *P* < .001; **Fig. S1F**). Excluding these three studies led to a data set without additional outliers (**Fig. S2**), which was used for the subsequent analyses.

As expected from the diversity of covariates such as behavioral outcome measures and rehabilitation types across studies, heterogeneity remained substantial (thoracic: *Q*_*E*_ (215) = 570.13, *P* < 0.001; cervical: *Q*_*E*_ (72) = 122.22, *P* < 0.001; **Fig. S2F**). The pooled effect size estimates revealed strong subgroup differences (*Q*_*M*_ (1) = 19.17, *P* < 0.001) between rehabilitation outcomes of thoracic and cervical injuries. Animals with thoracic spinal cord injuries showed approximately twice the SMD of rehabilitation intervention compared to those with cervical injuries (0.854, 95% CI: 0.715–0.992 vs. 0.466, 95% CI: 0.363–0.570). These findings support stratifying analyses by spinal level, as the observed differences in rehabilitation efficacy likely reflect fundamental distinctions in functional outcome measures, injury biology, and/or regenerative capacity between thoracic and cervical spinal cord injuries.

To account for the contribution of behavioral outcome measures to the heterogeneity, we stratified the multi-level meta-analysis accordingly. Stratification greatly reduced heterogeneity within the cervical dataset, although significant residual heterogeneity remained for forepaw ladder/grid performance and reaching success. BBB/BMS scores and hindpaw ladder/grid outcomes showed no significant residual heterogeneity (**Table S2, Fig. S3**). For the thoracic dataset, heterogeneity persisted for several behavioral measures, including the Basso, Beattie and Bresnahan (BBB) locomotor rating scale / Basso mouse scale (BMS), hindpaw ladder/grid, coordination, swim score, and stride length (**Table S3, Fig. S4**). This residual heterogeneity likely reflects the influence of the different rehabilitation types. To better account for these sources of variability, we stratified analyses by spinal level and behavioral measure, and then subgrouped by rehabilitation type.

Behavioral outcomes with at least two supporting studies for cervical injuries were limited to forepaw and hindpaw ladder/grid scores, reaching success, and BBB/BMS scores (**Fig. 3, Table S2**). All outcomes showed moderate improvements attributed to rehabilitation, with SMD values ranging from 0.35 to 0.57, indicating a medium effect size (**Fig. 3**). Specifically, most improved behaviors were reaching success (0.57) and forepaw ladder/grid scores (0.51), followed by hindpaw ladder/grid scores (0.39) and BBB/BMS scores (0.35).

**Fig. 3.**
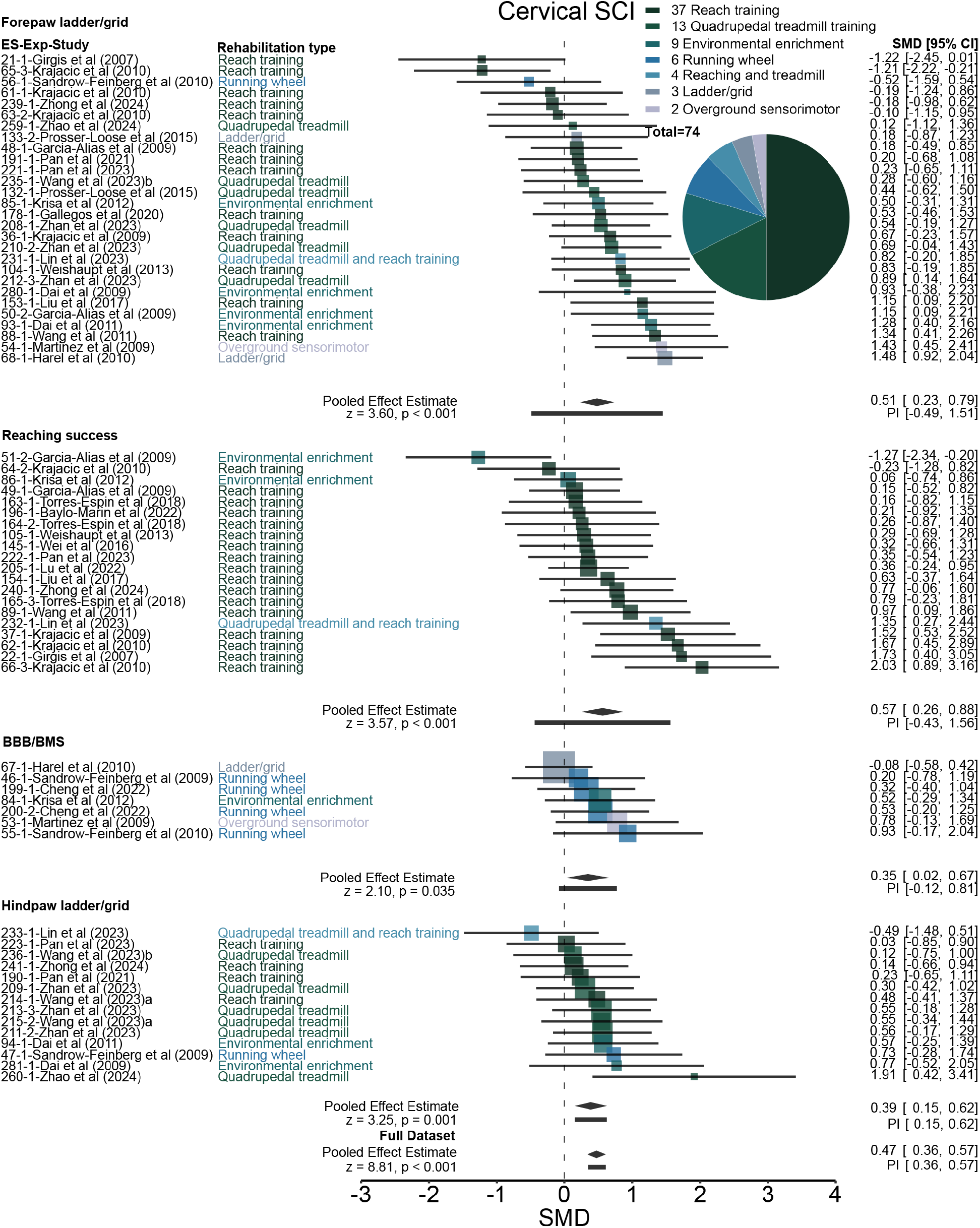
The effects of rehabilitation on behavioral outcomes following cervical-level spinal cord injuries. Each forest plot presents SMDs with 95% confidence intervals for individual experiments grouped by behavioral measure. The left-hand column lists the effect-size number (ES no.), experiment identifier (Exp), study name, and rehabilitation type, while the right-hand column contains the corresponding effect size and its associated 95% confidence interval. Rehabilitation types are color-coded for clarity. The size of each square reflects the weight assigned to the experiment by the random-effects multilevel model—larger squares indicate greater influence on the overall effect size. At the bottom of each behavioral measure, the total pooled effect estimates

Cervical-level injuries appeared largely more resistant to rehabilitation-induced improvements compared to thoracic-level injuries. Specifically, thoracic injury studies showed strong and consistent behavioral improvements across outcome measures in the model (**Figs. 4 and 5**). We found that rehabilitation after thoracic injury improved the BBB/BMS scores by an average of more than double compared to after a cervical injury (SMD of 0.83 in thoracic injuries vs. 0.35 in cervical injuries) (**Figs. 3 and 4; Tables S2 and S3**). Similarly, the hindpaw ladder/grid score showed a more than 2-fold higher improvement in thoracic injuries as compared to cervical injuries (SMD of 0.90 in thoracic injuries vs. 0.39 in cervical injuries, (**Figs. 3 and 5**). This suggests that injury level may be a critical determinant of functional recovery gains following rehabilitation in rodent spinal cord injury models.

**Fig. 4.**
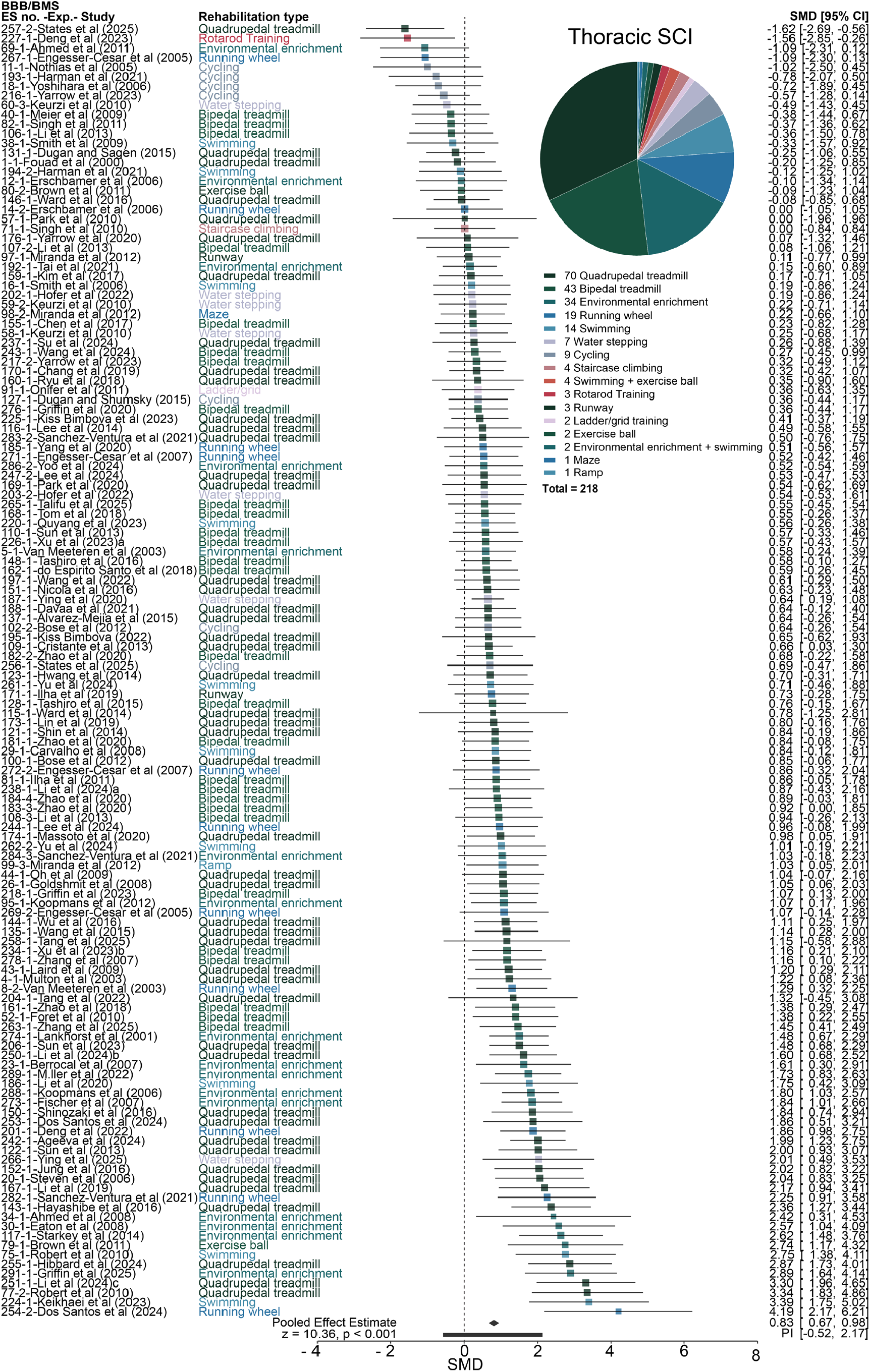
The effects of rehabilitation on BBB/BMS scores following thoracic-level spinal cord injuries. Forest plot presents SMDs with 95% confidence intervals for individual experiments measuring BBB/BMS scores. The left-hand column lists the effect-size number (ES no.), experiment identifier (Exp), study name, and rehabilitation type, while the right-hand column contains the corresponding effect size and its associated 95% confidence interval. Rehabilitation types are color-coded for clarity. The size of each square reflects the weight assigned to the experiment by the random-effects multilevel model—larger squares indicate greater influence on the overall effect size. The total pooled effect estimate is reported at the bottom of the forest plot.

**Fig. 5.**
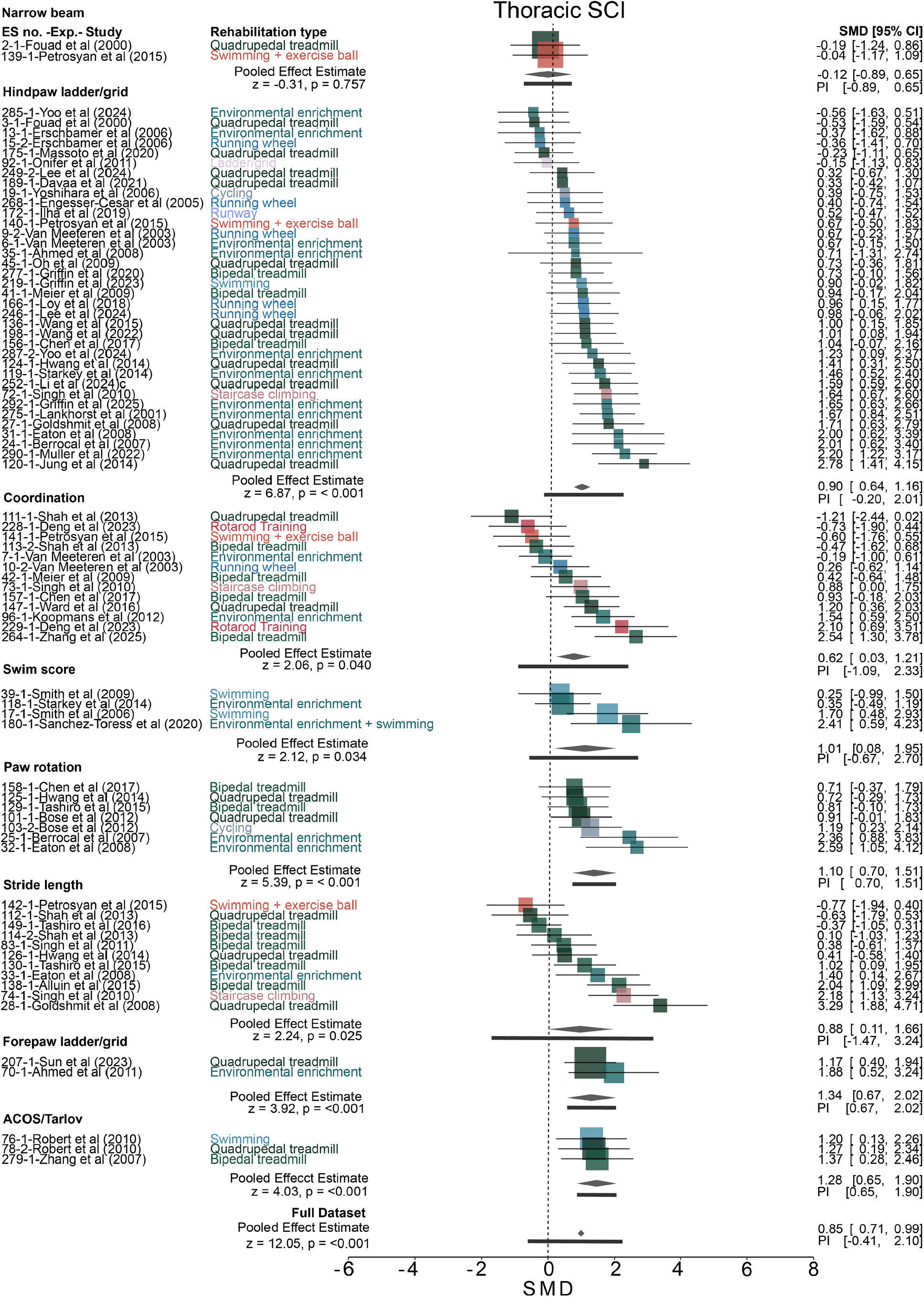
The effects of rehabilitation on behavioral outcomes following thoracic-level spinal cord injuries. Forest plot presents SMDs with 95% confidence intervals for individual experiments grouped by behavior measures. The left-hand column lists the effect-size number (ES no.), experiment identifier (Exp), study name, and rehabilitation type, while the right-hand column contains the corresponding effect size and its associated 95% confidence interval. Rehabilitation types are color-coded for clarity. The size of each square reflects the weight assigned to the experiment by the random-effects multilevel model—larger squares indicate greater influence on the overall effect size. At the bottom of each behavioral measure, the total pooled effect and the model-derived effect estimate are reported.

### Environmental enrichment is a key driver of functional recovery after experimental SCI

Across injury levels and behavioral outcomes, we noted that studies employing environmental enrichment–based rehabilitation tended to yield larger SMDs relative to other intervention strategies (**Figs. 3-5**). To investigate how this study-level characteristic (moderator) influences the overall effect, we employed meta-regression with categorical moderation (group comparison) and report the meta-regression coefficients (**Fig. 2C**). This analysis allowed direct comparison of rehabilitation strategies to determine whether specific approaches conferred greater functional benefits over others.

Consistent with our earlier analyses, overall effect sizes remained modest for cervical injuries (**Fig. 6A, Table S4**) compared to thoracic injuries (**Fig. 6B, Table S5**). In cervical SCI models, forepaw function assessed via the error ladder or grid test showed the largest improvements with environmental enrichment (SMD = 0.971 ± 0.315, *P* = 0.002) and ladder/grid training (SMD = 0.991 ± 0.391, *P* = 0.011). In contrast, we found that quadrupedal treadmill training (SMD = 0.524 ± 0.2929, *P* = 0.074) and reach training (SMD = 0.257 ± 0.181, *P* = 0.156) did not significantly improve forepaw performance (**Fig. 6A, Table S4**). For hindpaw ladder/grid scores, among reach training, environmental enrichment, and quadrupedal treadmill training, only the treadmill training yielded improvements (SMD = 0.503 ± 0.173, *P* = 0.004). BBB/BMS scores and reaching success were less frequently assessed. Nevertheless, running wheel rehabilitation moderately improved BBB/BMS scores (SMD = 0.457 ± 0.214, *P* = 0.033), and reach training enhanced reaching success (SMD = 0.638 ± 0.146, *P* < 0.001).

**Fig. 6.**
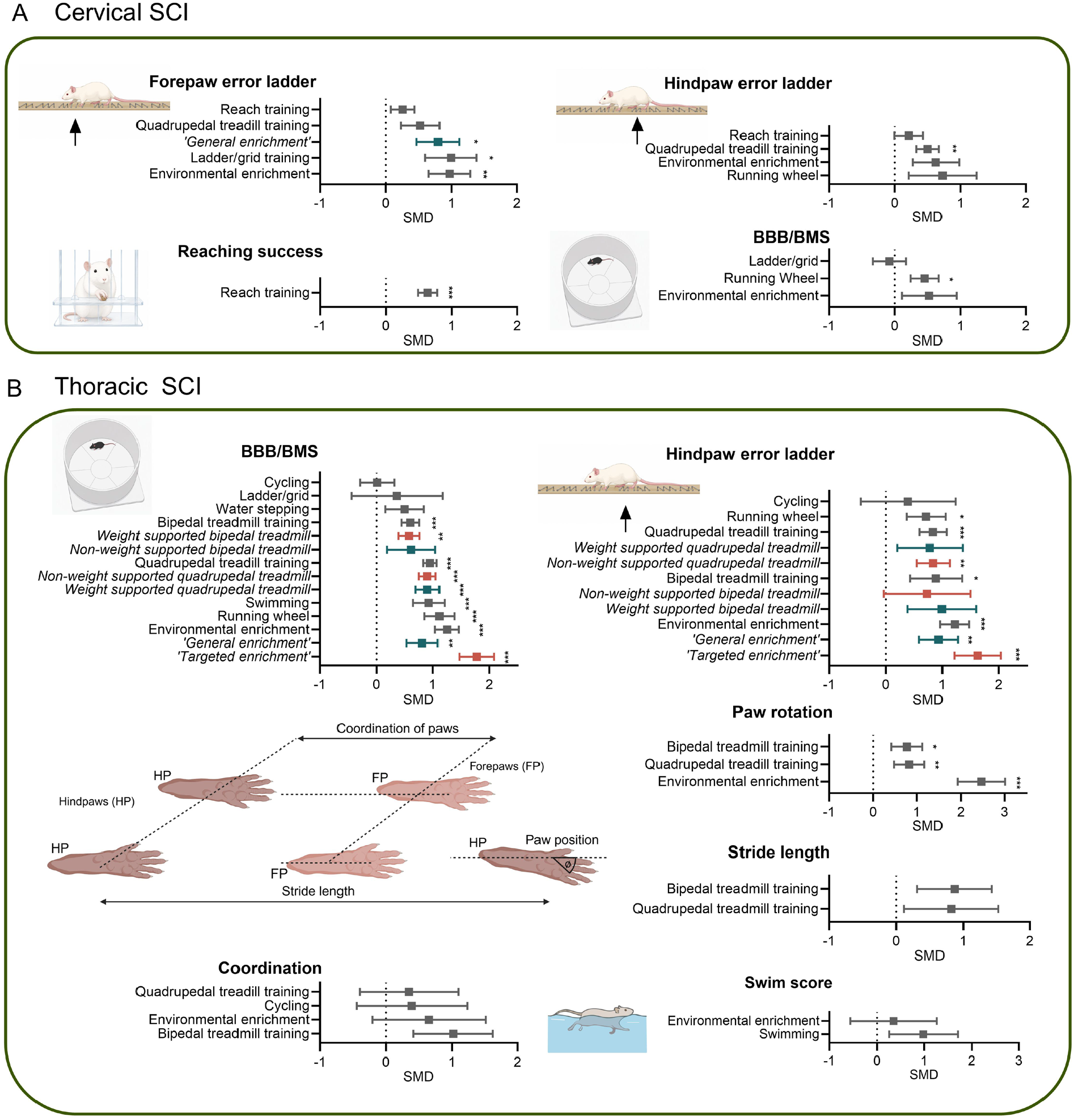
Rehabilitation efficacy varies according to injury level, behavioral outcome, and type of rehabilitation. **(A)** Categorical meta-regression of rehabilitation interventions in cervical SCI models. Effect sizes (SMD) are shown for forepaw error ladder, hindpaw error ladder, reaching success, and BBB/BMS score outcomes. Environmental enrichment and task-specific reaching paradigms generally produced the largest improvements in forelimb and skilled motor outcomes. **(B)** Categorical meta-regression of rehabilitation interventions in thoracic SCI models. Outcomes are stratified by behavioral assessment, including BBB/BMS scores, hindpaw error ladder performance, gait parameters (stride length, paw rotation, and coordination), and swim score. Data are presented as SMD with 95% confidence intervals. Positive values indicate improved behavioral performance following rehabilitation relative to control animals. Colors denote rehabilitation modality subcategories. Asterisks indicate statistical significance of the pooled effect estimate (^*\**^*P* < 0.05, ^**^*P* < 0.01, ^***^*P* < 0.001).

For thoracic-level injuries (**Fig. 6B, Table S5**), the larger dataset enabled more comprehensive comparisons. In fact, environmental enrichment produced robust improvements across multiple measures: BBB/BMS scores (SMD = 1.247 ± 0.208, *P* < 0.001), hindpaw ladder/grid scores (SMD = 1.218 ± 0.255, *P* < 0.001), and paw rotation (SMD = 2.467 ± 0.542, *P* < 0.001). Importantly, further analysis of rehabilitation subcategories revealed that targeted environmental enrichment, which incorporates task-specific training elements, approximately doubled the improvement in BBB/BMS scores compared to general environmental enrichment (SMD = 1.775 vs. 0.809, **Table S6**) and produced greater improvements in hindpaw ladder/grid performance (SMD = 1.623 vs. 0.932; **Table S6**). This suggests that while general environmental enrichment helps motor recovery, task-specific targeting further enhances functional recovery.

For hindpaw function on the error ladder/grid test, both bipedal (SMD = 0.890 ± 0.459) and quadrupedal treadmill training (SMD = 0.836 ± 0.238, *P* < 0.001) yielded positive effect estimates, although the bipedal treadmill estimate narrowly missed statistical significance (*P* = 0.053). For hindpaw ladder/grid performance, running-wheel exercise produced the smallest effect estimate among these interventions, although the improvement remained significant (SMD = 0.714 ± 0.342, *P* = 0.037). Quadrupedal and bipedal treadmill training yielded comparable improvements in hindpaw ladder/grid scores and paw rotation. However, quadrupedal treadmill training produced a larger estimated improvement in BBB/BMS scores (SMD = 0.952 ± 0.119, *P* < 0.001) than bipedal treadmill training (SMD = 0.601 ± 0.157, *P* < 0.001, **Table S5**). Notably, no significant differences were observed between quadrupedal treadmill training with or without weight support (**Table S6**).

Although the behavior-stratified model indicated an overall improvement in coordination, no individual rehabilitation modality produced a statistically significant coordination effect in the rehabilitation-type meta-regression. Swimming training failed to improve swimming scores, nor did environmental enrichment significantly affect swimming performance. However, swimming rehabilitation led to substantial improvements in BBB/BMS scores (SMD = 0.928 ± 0.279, P < 0.001). Water-stepping and cycling generally produced smaller or less consistent improvements than environmental enrichment and treadmill-based rehabilitation.

In summary, environmental enrichment consistently produced the most pronounced functional recovery across behavioral measures, particularly in thoracic SCI models. Targeted environmental enrichment outperformed general environmental enrichment, while more aversive methods such as cycling and water-stepping were the least beneficial. These findings prompted further investigation into how other study-level characteristics affect rehabilitation outcomes.

### Longer rehabilitation duration is associated with greater recovery, with no detectable plateau within the available data

Longer inpatient rehabilitation improves functional outcomes for patients with SCI. Specifically, studies using the motor Functional Independence Measure (FIM) show that longer rehabilitation stays are associated with greater functional gains between admission and discharge (*33-35*). However, evidence demonstrates a clear plateau effect where spontaneous functional recovery slows substantially. Driven by biological recovery timelines and motor learning constraints, most of the neurological recovery occurs within the first 3–6 months after injury, with recovery rates declining thereafter and approaching a plateau by 9–12 months (*36-38*). We therefore investigated whether a similar duration-response relationship exists in preclinical rehabilitation studies and whether treatment benefits exhibit evidence of plateauing with increasing intervention duration.

To assess the relationship between rehabilitation duration and treatment efficacy, we performed a mixed-effects meta-regression with total intervention duration (in weeks total) as a moderator and rehabilitation type and behavioral outcome measure included as random effects (**Fig. 2**). The moderator model was significant (*Q*_*M*_(1) = 4.51, *P* = 0.034; **Table S7**), indicating that intervention duration explained a significant proportion of the variability in treatment effects. A positive association was observed between intervention duration and efficacy (β = 0.02270 ± 0.0106, 95% CI 0.0017–0.0436, P = 0.034; **Fig. 7A, Table S7**), corresponding to an increase of approximately 0.023 SMD units for each additional week of rehabilitation. The intercept was 0.6710 (95% CI 0.435–0.906, *P* < 0.001). Although significant residual heterogeneity remained (*Q*_*E*_(209) = 558.33, *P* < 0.001), these findings suggest that longer rehabilitation paradigms are associated with greater functional recovery.

**Fig. 7.**
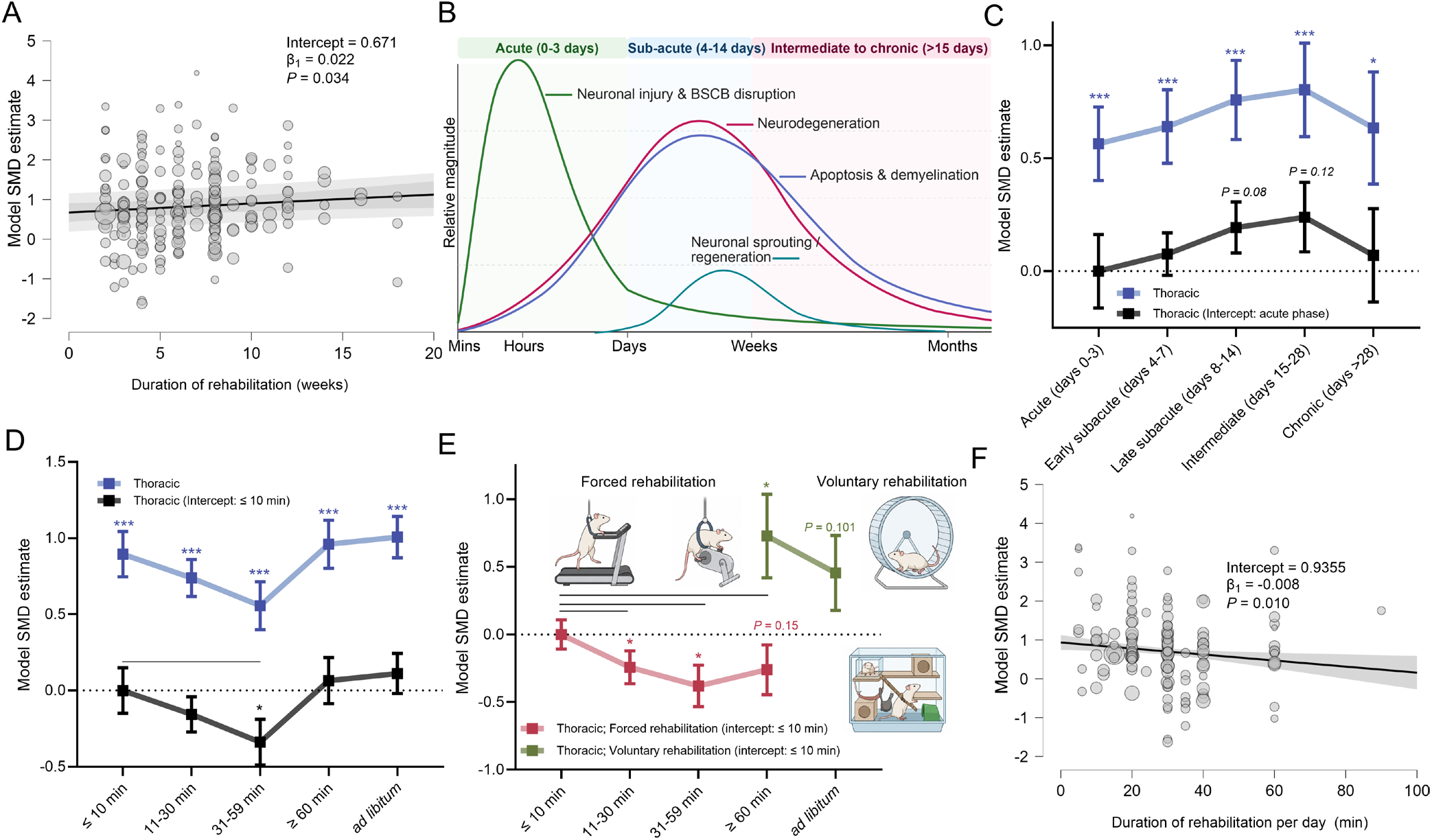
Rehabilitation duration, timing, and delivery mode influence functional recovery following spinal cord injury. **(A)** Mixed-effects meta-regression demonstrating a positive association between total rehabilitation duration (weeks) and treatment efficacy. Longer rehabilitation interventions were associated with greater improvements in functional outcomes. (**B**) Schematic representation of key biological processes occurring across the acute, subacute, and chronic phases following spinal cord injury, illustrating the temporal evolution of tissue damage, neurodegeneration, and regenerative plasticity. **(C)** Effect of rehabilitation onset phase on treatment efficacy. Blue symbols represent estimated effect sizes within each injury phase from a no-intercept model, whereas black symbols represent comparisons relative to the acute phase from an intercept model. Rehabilitation produced significant functional improvements across all injury stages, with no significant differences observed between phases. **(D)** Effect of daily rehabilitation exposure on treatment efficacy. Blue symbols represent pooled effect sizes within each rehabilitation-duration category, whereas black symbols indicate differences relative to the reference category (≤10 min/day). Treatment efficacy varied according to daily rehabilitation exposure, with the 31–59 min/day category demonstrating significantly smaller effects than the shortest-duration category. **(E)** Subgroup analysis examining the interaction between rehabilitation duration and rehabilitation modality. Increasing duration was associated with reduced efficacy in forced rehabilitation paradigms, whereas voluntary rehabilitation tended to benefit from prolonged access, with interventions exceeding 60 min/day producing significantly larger effects than the shortest-duration category. **(F)** Continuous meta-regression of forced rehabilitation duration demonstrating a significant negative relationship between daily rehabilitation exposure and treatment efficacy, indicating that prolonged forced rehabilitation was associated with diminished functional gains. Data are presented as SMDs and 95% confidence intervals. Asterisks indicate significant differences from the reference category (^*\**^*P* < 0.05, ^*\**\*^*P* < 0.01, ^*\**\*\*^*P* < 0.001).

To determine whether this duration-response relationship reached a plateau, we next fitted a quadratic meta-regression incorporating both linear and squared duration terms. Although the overall model approached significance (*Q*_*M*_(2) = 5.37, *P* = 0.068; **Table S8**), neither the linear term (β = 0.056, 95% CI −0.016 to 0.129, *P* = 0.129) nor the quadratic term (β = −0.0021, 95% CI −0.0065 to 0.0023, *P* = 0.343) was significant. While the negative quadratic coefficient is directionally consistent with diminishing returns at longer durations, the effect was small and not statistically supported. Consequently, we found no evidence that rehabilitation efficacy plateaued within the relatively short treatment durations examined in the preclinical literature.

Collectively, these findings suggest that extending rehabilitation duration is associated with greater functional improvement in preclinical SCI models and, importantly, provide no statistical evidence that treatment benefits plateau within the intervention timeframes currently employed. This contrasts with the temporal constraints observed for spontaneous neurological recovery and supports the possibility that rehabilitation-driven functional gains remain responsive to continued training beyond periods typically associated with endogenous recovery processes.

### A broad therapeutic window supports rehabilitation efficacy across the continuum of spinal cord injury

The optimal timing of rehabilitation after SCI remains a topic of ongoing debate. Proponents of early intervention argue that rehabilitation should begin during a critical period of heightened neuroplasticity, as delaying treatment for weeks or months may substantially reduce the potential for functional recovery (*39*). In contrast, others contend that rehabilitation initiated too soon after injury may coincide with acute pathological processes that limit its effectiveness or even exacerbate tissue damage (*40, 41*). These competing perspectives may reflect the dynamic cascade of biological events that follows SCI (**Fig. 7B**), whereby the acute post-injury period is dominated by inflammation and secondary injury mechanisms, while subacute phases are characterized by increased opportunities for neural plasticity and regeneration. Consequently, rehabilitation initiated too early may be hindered by an unstable microenvironment, whereas delayed intervention may miss critical windows of neuroplastic potential. As a result, a clear consensus regarding the optimal therapeutic window has yet to emerge, and the timing of rehabilitation is likely to depend on both the intervention modality and the underlying injury model.

To determine whether the timing of rehabilitation influences treatment efficacy, injury onset phase was evaluated as a categorical moderator in a mixed-effects meta-regression, with rehabilitation type and behavioral outcome measure included as random effects. Injury phases were classified as acute (days 0-3), early subacute (days 4-7), late subacute (days 8-14), intermediate (days 15-28), or chronic (>28 days). Across studies, rehabilitation produced a significant improvement in functional outcomes (SMD = 0.65, 95% CI 0.34–0.95, *P* < 0.001, **Table S9**).

Using an intercept model with acute injury as the reference category, onset phase did not significantly moderate treatment efficacy (*Q*_*M*_(4) = 4.34, *P* = 0.362, **Table S9**). Relative to acute injury, treatment effects were not significantly different during the early subacute (β = 0.08, 95% CI −0.11 to 0.26, *P* = 0.420, **Fig. 7C**), late subacute (β = 0.19, 95% CI −0.03 to 0.42, *P* = 0.087), intermediate (β = 0.24, 95% CI −0.06 to 0.54, *P* = 0.120), or chronic phases (β = 0.07, 95% CI −0.33 to 0.47, *P* = 0.735), indicating that rehabilitation efficacy was largely preserved across the injury continuum.

To estimate treatment efficacy within each injury phase in greater detail, a no-intercept model was subsequently fitted. Under this parameterization, rehabilitation was associated with significant functional improvement regardless of when treatment was initiated, with effect sizes ranging from 0.56 in the acute phase (95% CI 0.25–0.88, *P* < 0.001, **Fig. 7C, Table S9**) to 0.80 in the intermediate phase (95% CI 0.40–1.21, *P* < 0.001). Numerically larger effects were observed during the late subacute (SMD = 0.76, 95% CI 0.41–1.10, *P* < 0.001) and intermediate phases, whereas acute (SMD = 0.56, 95% CI 0.25–0.88, *P* < 0.001) and chronic injury (SMD = 0.63, 95% CI 0.15–1.12, *P* = 0.011) exhibited more modest improvements.

Collectively, these findings indicate that rehabilitation remains effective across a broad therapeutic window following SCI in rodents, with no strong evidence that treatment efficacy is restricted to a specific stage of injury. Although effect sizes were numerically greater during the late subacute and intermediate phases, the absence of significant between-phase differences suggests that the capacity for rehabilitation-driven functional improvement is maintained throughout both subacute and chronic stages of recovery. This broad window of responsiveness is consistent with the persistence of activity-dependent neuroplasticity after SCI and supports rehabilitation as a viable intervention strategy across multiple timepoints of injury progression.

### Functional recovery varies according to daily rehabilitation exposure

While longer cumulative rehabilitation exposure is generally associated with improved functional recovery, increasing treatment duration alone may not be sufficient to maximize outcomes once recovery trajectories begin to plateau (*42*). Instead, the intensity and quality of rehabilitation are thought to become increasingly important drivers of recovery. Given the challenges associated with quantifying rehabilitation intensity across preclinical studies, we used daily rehabilitation duration as a pragmatic surrogate for rehabilitation dosage. We therefore performed a mixed-effects meta-regression examining whether treatment efficacy varied across five daily rehabilitation exposure categories (≤10 min, 11–30 min, 31–59 min, ≥60 min, and ad libitum access), while accounting for variation attributable to rehabilitation modality and behavioral outcome measure through random-effects modelling.

The overall moderator effect was significant (*Q*_*M*_(4) = 15.07, *P* = 0.005, **Table S10**), indicating that the daily rehabilitation exposure time explained a significant component of the variability in treatment effects. Using the shortest rehabilitation timeframe (≤ 10 min) as the reference category, studies in which rehabilitation was delivered for 31–59 min/day demonstrated significantly smaller treatment effects (β = –0.34, 95% CI –0.63 to –0.04, *P* = 0.024, **Fig. 7D, Table S10**). In contrast, medium-duration rehabilitation (11–30 min, β = –0.16, 95% CI –0.38 to 0.07, *P* = 0.176), or rehabilitation duration over 60 min (β = 0.06, 95% CI –0.23 to 0.36, *P* = 0.667), and ad libitum access to rehabilitation (β = 0.11, 95% CI –0.15 to 0.37, *P* = 0.396) did not differ significantly from the reference group.

To estimate efficacy within each category, a no-intercept model was subsequently fitted. This analysis revealed significant treatment effects across all rehabilitation timeframes, although effect sizes varied considerably between categories (**Fig. 7D, Table S10**). The largest effects were observed when rehabilitation occupied the shortest proportion of the study period (SMD = 0.90), whereas studies categorized as long-duration (31–59 min) rehabilitation demonstrated comparatively smaller effect sizes. Effect sizes increased again for rehabilitation durations of more than 60 min per day as well as ad libitum access, suggesting a non-linear relationship between rehabilitation allocation and functional recovery which may reflect underlying study characteristics.

Collectively, these findings indicate that the daily rehabilitation duration influences treatment efficacy. However, the observed pattern does not support a simple dose-response relationship, as extending rehabilitation over a greater proportion of the study period was not consistently associated with larger treatment effects. Instead, efficacy appeared to vary across rehabilitation schedules, suggesting that factors such as intervention intensity, timing, and study design may interact with rehabilitation duration to influence recovery outcomes.

### Forced and voluntary rehabilitation differ in their dose-response relationships

Because physiological stress is known to impair functional recovery following neurological injury (*43-46*), we hypothesized that stress associated with forced rehabilitation paradigms may contribute to the diminished efficacy observed with increasing training duration. Consistent with this hypothesis, treadmill training induces robust systemic stress responses in both male and female rodents (*47*), whereas voluntary wheel running and environmental enrichment are generally not associated with comparable elevations in physiological stress markers (*48*). We therefore examined whether the relationship between rehabilitation exposure and treatment efficacy differed between forced and voluntary rehabilitation paradigms.

Subgroup-specific mixed-effects meta-regressions were performed with daily rehabilitation duration entered as a categorical moderator and behavioral outcome measure and rehabilitation type entered as random effects. Within forced rehabilitation studies, the shortest-duration category (≤10 min/day) produced a large treatment effect (SMD = 0.94, 95% CI 0.73–1.15, *P* < 0.001, **Fig. 7E, Table S11**). Increasing rehabilitation exposure was associated with progressively smaller treatment effects, with both the 11–30 min/day (β = −0.24, 95% CI −0.48 to −0.00, *P* = 0.046) and 31–59 min/day categories (β = −0.38, 95% CI −0.68 to −0.08, *P* = 0.013) demonstrating significantly lower efficacy than the reference group. Rehabilitation durations exceeding 60 min/day showed a similar trend, although this did not reach significance (β = −0.26, 95% CI −0.62 to 0.10, *P* = 0.157).

In contrast, voluntary rehabilitation exhibited the opposite pattern. Relative to the shortest-duration category, interventions providing more than 60 min/day of voluntary activity were associated with significantly larger treatment effects (β = 0.73, 95% CI 0.12–1.33, *P* = 0.018, **Fig. 7E, Table S11**), while *ad libitum* access showed a similar but non-significant trend (β = 0.45, 95% CI −0.09 to 1.00, *P* = 0.101). Collectively, these findings suggest that prolonged rehabilitation exposure may be beneficial when activity is self-directed but not when it is externally imposed.

To determine whether this relationship was dose dependent, daily duration was subsequently modeled as a continuous moderator within forced rehabilitation studies. This analysis identified a negative association between rehabilitation duration and treatment efficacy (*Q*_*M*_(1) = 6.57, P = 0.010; **Fig. 7F, Table S12**), with each additional minute of rehabilitation associated with an approximate 0.008-unit reduction in effect size (β = −0.0078, 95% CI −0.0137 to −0.0018, *P* = 0.010). These findings indicate that increasing the duration of forced rehabilitation per day does not necessarily translate into greater functional gains and may instead reduce treatment efficacy.

To further explore the potential contribution of stress in this outcome, rehabilitation paradigms were assigned qualitative stress grades ranging from 0 to 3 according to their established propensity to elicit physiological stress responses (46,47; **Supplementary Methods**). Low-stress, voluntary interventions such as environmental enrichment were assigned a grade of 0, whereas highly demanding paradigms, including cycling and water stepping, were assigned a grade of 3. Consistent with our hypothesis, grade-0 interventions produced the largest treatment effects, whereas grade-3 interventions were associated with minimal functional improvement (**Fig. S5A**). We next integrated rehabilitation exposure and stress burden by calculating a composite stress coefficient (stress grade × daily rehabilitation duration; **Supplementary Methods)**. This coefficient was negatively correlated with treatment efficacy (r = −0.292, *P* < 0.001; **Fig. S5B**), suggesting that cumulative stress exposure may limit the benefits of rehabilitation after SCI.

Together, these findings indicate that rehabilitation efficacy depends not only on the amount of training delivered but also on how that training is experienced. Whereas prolonged forced rehabilitation was associated with reduced efficacy, voluntary approaches appeared to benefit from extended access. This pattern may help explain the superior outcomes observed with low-stress, self-directed interventions such as environmental enrichment relative to more demanding modalities, including cycling and water stepping. Notably, voluntary rehabilitation strategies, particularly environmental enrichment initiated during the late subacute phase, emerged as some of the most effective approaches for promoting recovery after SCI.

## DISCUSSION

Despite the widespread use of rehabilitation in preclinical SCI research, the factors that determine its efficacy remain poorly understood. In this comprehensive meta-analysis of 150 studies spanning more than 3,000 animals, 292 effect sizes, and 12 behavioral outcomes, we identified environmental enrichment as the most effective rehabilitation strategy for promoting functional recovery after SCI. These benefits were particularly pronounced in thoracic injury models, where effect sizes generally exceeded those observed following cervical injury. Although rehabilitation was effective across all injury stages, the largest numerical effect was observed during the intermediate phase, followed closely by the late subacute phase; neither differed significantly from the acute phase. We further found that increasing the duration of forced rehabilitation was associated with reduced therapeutic benefit, whereas voluntary approaches, such as environmental enrichment, appeared to benefit from prolonged exposure. This pattern was accompanied by a negative association between rehabilitation efficacy and our exploratory composite measure of stress burden, suggesting that stress may be an important, yet often overlooked, modifier of recovery. Collectively, these findings demonstrate that the effectiveness of rehabilitation depends not only on the amount of training delivered but also on when and how it is delivered, emphasizing the need to consider species-specific behavioral, anatomical, and physiological factors when designing and interpreting preclinical rehabilitation studies. Before discussing these findings in detail, it is important to consider the key differences between recovery in rodents and humans following SCI.

### Differences between the human and rodent recovery profiles

Indeed, a notable divergence emerges between human and rodent recovery profiles following SCI. Clinically, individuals with cervical injuries are more likely to improve by at least one American Spinal Injury Association Impairment Scale grade than those with thoracic injuries, with reported improvement rates of approximately 37% for cervical versus 16% for thoracic injuries (*49-53*). In contrast, our meta-analysis of preclinical data revealed greater-standardized functional recovery following thoracic compared to cervical injuries in rodents. This apparent discrepancy likely reflects fundamental anatomical and behavioral differences between species that shape recovery trajectories. From an anatomical perspective, spinal tracts critical for motor recovery—particularly those involved in relay network formation such as the reticulospinal tract—are positioned more medially in the human spinal cord, rendering them especially vulnerable to contusion injury. In rodents, these tracts are located more laterally and are therefore more likely to be spared following injury. Even minimal preservation of reticulospinal fibers in rodents is strongly associated with improved motor recovery (*54*), in part through the formation of novel synaptic connections with propriospinal neurons that bypass the primary lesion site (*55, 56*). This capacity for circuit reorganization may underlie the more pronounced recovery observed in rodent models. Reorganization is further enhanced by neuronal activity, which modulates axon growth, synapse formation, and transmission—processes increasingly recognized as central to rehabilitation-driven plasticity (*57, 58*). Notably, axon growth and synaptic stabilization may be partially competing processes. Consistent with this view, electrical silencing or hampering synaptic transmission can enhance axon regeneration of some neurons, including dorsal root ganglion neurons (*3, 59, 60*). Hence, electrical activity and rehabilitation may be essential for shaping new connections, not for long-distance regeneration *per se*. Thus, there are clear differences between human and rodent recovery. Taking these differences into account helps to adjust translational parameters.

### Does task-specific training come at a cost?

Differences in recovery also occur because humans are bipedal and rodents are quadrupedal. Human rehabilitation, while multifactorial addressing many aspects of recovery, functionally often prioritizes ambulatory upper limb function, potentially limiting engagement of lower limb motor circuits (*61*). In contrast, in rodents, whole-body movement is either intentionally or inadvertently encouraged through innate rodent behavior, housing conditions and/or experimental conditions. This is likely one reason that rodents spontaneously recover even after severe injury, particularly in hindlimb function, where animals can progress from complete paralysis to weight-supported plantar stepping in a matter of two weeks. - Therefore, ‘off-time’ factors are important to consider in animal models. Consistent with this, experimental models demonstrate that limb disuse leads to functional decline: rodents subjected to forelimb immobilization after SCI or stroke exhibit pronounced deficits in the immobilized limb (*62, 63*), a finding replicated across multiple paradigms. Together, these anatomical and behavioral considerations underscore the central role of activity-dependent plasticity in rodent recovery and highlight important limitations in directly extrapolating preclinical rehabilitation outcomes to the clinical setting (*64-66*).

A long-standing debate in neurorehabilitation concerns the potential trade-offs of general and task-specific training (*67*). Early studies in spinalized cats demonstrated that stand-training improved standing ability but impaired overground walking, and vice versa (*68*). Similar findings have been reported in rodents: general environmental enrichment was shown to diminish skilled paw function (*69*), and skilled reach training impaired performance on the ladder task (*70*). More recently, however, it was shown that standing and locomotor recovery in cats is due primarily to the return of neuronal excitability within spinal sensorimotor circuits independent of task-specific activity-dependent plasticity (*71*). Our analysis provides new evidence relevant to this debate, raising the question: is the decline in those studies simply due to neglect of untrained functions? – similar to findings from limb immobilization studies. For environmental enrichment-based rehabilitation, animals are exposed to a broad array of movements and stimuli, potentially mitigating the risk of such trade-offs by promoting more generalized recovery. In fact, a targeted environmental enrichment approach can be designed to actively promote functional recovery through deliberate inclusion of rehabilitative tasks aimed at promoting diverse movement patterns directed at restoring lost function (*72*).

### Environmental enrichment

The benefits of environmental enrichment are believed to arise from a convergence of factors, including enhanced social interaction, improved mood, and the integration of diverse therapeutic modalities (*73*). At the neurobiological level, enrichment has been associated with a range of adaptive processes, such as synaptogenesis, dendritic arborization, axonal sprouting, neurogenesis, synaptic plasticity, neuronal survival, and angiogenesis (*11*). Increased sensory stimulation also plays a pivotal role in driving these changes (*74, 75*). Collectively, these findings support the notion that broad, integrative approaches such as environmental enrichment may be particularly effective through multiple factors. It may also be important to avoid an overly neurocentric interpretation of these mechanisms. While neural plasticity is fundamental to recovery, systemic physiological processes such as vascular health and musculoskeletal integrity are equally vital for functional ability in rodents. Sustained physical activity is crucial for preventing muscle atrophy and maintaining cardiovascular function in clinical populations (*76-78*). Therefore, the therapeutic benefits of enrichment likely emerge from the dynamic interplay between neural, vascular, and systemic adaptations.

### Total duration and cumulative gains

Our analysis indicates that a longer total rehabilitation duration is associated with larger treatment effects, supporting the hypothesis that sustained training enhances functional recovery after SCI. In contrast to the well-described plateau in spontaneous neurological recovery observed clinically, we found no statistical evidence that rehabilitation efficacy plateaued within the intervention durations evaluated. Although the quadratic model yielded a negative coefficient, suggesting a potential trend toward diminishing returns at longer durations, this effect was not significant and therefore provides insufficient evidence for a true ceiling effect. One interpretation is that rehabilitation-driven recovery in rodents reflects ongoing experience-dependent neuroplasticity that remains responsive to continued practice beyond the period during which spontaneous recovery predominates. Alternatively, and more likely, the absence of a detectable plateau may reflect the limited number of studies employing long-duration rehabilitation protocols – the average rehabilitation duration was four to six weeks. These findings suggest that extending treatment duration may represent a straightforward strategy for maximizing functional recovery in preclinical SCI studies and reinforce the need for future studies specifically designed to define the optimal dose and duration of rehabilitation.

### Timing of rehabilitation initiation

The timing of when to initiate rehabilitation interventions following SCI also remains a subject of ongoing debate, with no clear consensus emerging from either clinical or preclinical research. Despite decades of investigation, we still lack definitive guidelines for when and how intensively to initiate rehabilitation in both human and rodent models. While a recent clinical trial aimed at addressing the question of onset has been pre-registered (NCT06176833; (*79*)), robust clinical evidence remains limited. Preclinical studies have suggested a delicate balance may be at play: initiating rehabilitation too early may provoke maladaptive neuroplasticity, while delaying intervention may miss the critical window of heightened plastic potential following injury (*40, 41*).

Contrary to the expectation that rehabilitation efficacy may be restricted to a narrow post-injury window, our meta-regression found no significant differences in treatment efficacy across acute, subacute, intermediate, and chronic injury phases. Although effect sizes were numerically greatest during the late subacute and intermediate phases, these differences were not statistically significant when compared to the acute phase. Importantly, rehabilitation produced significant benefits within all injury stages examined, indicating that the capacity for rehabilitation-driven functional improvement is maintained throughout the course of injury.

These findings suggest that activity-dependent plasticity remains accessible well beyond the acute phase of injury and may support meaningful recovery even after spontaneous neurological recovery has begun to slow. This observation is consistent with experimental and clinical evidence demonstrating persistent remodeling of spared neural circuits, adaptive changes within spinal and supraspinal networks, and ongoing capacity for motor learning during chronic SCI. From a translational perspective, these data challenge the perception that rehabilitation must be initiated within a narrowly defined therapeutic window to be effective and instead support the implementation of rehabilitation strategies across a broad range of injury stages. Such findings are particularly relevant for clinical populations, where access to intensive rehabilitation is often delayed and many individuals enter rehabilitation months or years after injury.

### Rehabilitation dose and delivery

A major finding of this meta-analysis was that rehabilitation efficacy depended not only on the amount of rehabilitation delivered, but also on how rehabilitation was delivered to rodents. Although greater rehabilitation exposure is generally assumed to promote recovery after SCI, we observed fundamentally different dose-response relationships for forced and voluntary rehabilitation paradigms. Increasing the duration of forced rehabilitation was associated with progressively smaller treatment effects, a pattern that was evident both when rehabilitation duration was categorized and when modeled as a continuous variable. In contrast, prolonged access to voluntary rehabilitation was associated with greater functional gains. These findings suggest that rehabilitation dose cannot be considered independently of the behavioral and physiological context in which training occurs and challenge the assumption that increasing training exposure alone is sufficient to maximize recovery.

One potential explanation for these findings is the contribution of physiological stress. Forced rehabilitation paradigms, including treadmill training, cycling, and water stepping, often require repeated handling, external motivation, or aversive stimuli, all of which may increase stress, fatigue, and task disengagement. By contrast, voluntary interventions such as environmental enrichment and wheel running allow animals to self-regulate training intensity and frequency while maintaining engagement. Consistent with this interpretation, interventions assigned to the highest stress grades demonstrated the smallest treatment effects, whereas low-stress interventions produced the greatest functional improvements. Moreover, the negative correlation between the composite stress coefficient and rehabilitation efficacy suggests that cumulative stress exposure may be an important factor limiting recovery after SCI. Elevated stress and corticosterone signaling have previously been linked to impaired behavioral outcomes and reduced neuroplasticity following SCI in rodents (*43-48*) with emerging evidence suggesting a similar relationship in humans (*80*).

The superior performance of voluntary rehabilitation paradigms may also reflect their capacity to engage motivation-dependent motor learning and activity-dependent plasticity. Self-directed activity allows animals to interact with their environment according to their functional abilities, potentially maximizing practice while minimizing fatigue. Environmental enrichment may additionally recruit reward-related neural circuitry, including nucleus accumbens signaling, to reinforce behavioral engagement and enhance recovery (*81*). This interpretation is consistent with the strong efficacy observed for environmental enrichment, particularly when initiated during the late subacute phase, when acute pathological processes have stabilized but the nervous system retains substantial capacity for adaptive reorganization. Environmental enrichment may therefore provide a biologically favorable combination of sustained activity, behavioral engagement, and neuroplastic stimulation that is not readily achieved through externally imposed exercise paradigms.

Importantly, these findings align with emerging clinical evidence suggesting that increasing rehabilitation intensity beyond conventional levels does not necessarily yield proportional improvements in outcome. The SCI-MT trial reported that adding intensive task-specific and strength-training interventions to standard inpatient rehabilitation failed to improve primary or secondary outcomes, highlighting the possibility of diminishing returns with increasing rehabilitation dose (*82-84*). Although direct translation between rodent and human rehabilitation should be approached cautiously, both the clinical and preclinical data suggest that there may be an upper threshold beyond which simply increasing rehabilitation exposure provides limited additional benefit. In rodents, excessive forced training may even impair recovery through stress-mediated mechanisms.

Collectively, these findings suggest that the quality, tolerability, and behavioral context of rehabilitation may be as important as its duration or intensity. Interventions that promote sustained, voluntary engagement appear to provide greater therapeutic benefit than more demanding rehabilitation paradigms and may better exploit neuroplastic mechanisms underlying recovery. Notably, recent clinical studies have begun translating principles of environmental enrichment into neurorehabilitation settings, demonstrating that structured environments promoting physical, cognitive, and social engagement can improve outcomes following stroke and non-progressive brain injury (*85, 86*). The convergence of these clinical observations with our preclinical findings supports the translational potential of environmental enrichment-inspired rehabilitation strategies and suggests that promoting engagement, autonomy, and multimodal activity may represent a promising framework for optimizing recovery after SCI.

### Conclusion

The substantial heterogeneity in preclinical spinal cord injury rehabilitation protocols, injury models, and treatment schedules provided an opportunity to identify factors associated with functional recovery. Together with improved data reporting across the field (*87*), this enabled systematic evaluation of the available evidence through meta-analysis. Our findings identify rehabilitation modality, duration, and delivery context as important determinants of efficacy, while supporting a broad therapeutic window for intervention.

Future preclinical studies should prioritize sustained voluntary engagement and incorporate task-specific, environmentally enriched activities where appropriate. Forced rehabilitation should be delivered cautiously to avoid excessive physiological stress, fatigue, or disengagement. Studies should distinguish between daily exposure, session frequency, total rehabilitation duration, and cumulative training dose, and should evaluate rehabilitation across a range of initiation times rather than assuming a narrow therapeutic window.

To improve reproducibility, investigators should clearly report rehabilitation onset, modality, task specificity, voluntary participation, weight support, session frequency and duration, progression criteria, and indicators of stress or adverse effects. Studies should also use appropriate injury-matched controls, validated behavioral outcomes, prespecified protocols, and blinded outcome assessment. These recommendations support a shift from defining rehabilitation primarily by training quantity toward interventions that promote sustained, task-relevant, and low-stress engagement. Their adoption will improve the rigor, reproducibility, and translational relevance of preclinical spinal cord injury rehabilitation research. Pairing candidate regenerative treatment with such optimized rehabilitation will help to more objectively extrapolate potential benefits for such treatment when translated to patients.

## MATERIALS AND METHODS

### Study design

The aim of this study was to conduct a comprehensive meta-analysis and meta-regression of peer-reviewed, published studies investigating rehabilitation modalities in preclinical models of SCI. We sought to address persistent gaps in understanding the comparative effectiveness of different rehabilitative interventions and the factors influencing functional recovery. To this end, we systematically screened a curated database for eligible studies, extracted detailed information on experimental designs and behavioral outcomes, and applied meta-analytical statistical techniques to quantify overall effect sizes, perform subgroup analyses, and conduct meta-regression. We began by comparing the overall effect sizes of rehabilitation interventions across cervical and thoracic injury datasets, using descriptive statistics to identify potentially influential observations. Multilevel random-effects meta-analysis models were then fitted using restricted maximum likelihood estimation while accounting for dependence among effect sizes and variability across behavioral outcome measures. Finally, meta-regression analyses were performed to evaluate the influence of rehabilitation characteristics, including rehabilitation type, onset phase, injury severity, and intervention duration, on functional outcomes.

### Defining the search strategy and eligibility criteria

A systematic literature search was conducted using the PubMed database with the following search terms: “*((spinal cord injury) OR (spinal cord injured) AND ((rehabilitation[Title]) OR (exercise[Title]) OR (training[Title]) OR (environmental enrichment[Title]) OR (enrichment[Title]) OR (swimming[Title]) OR (task-specific[Title]) OR (cycling[Title])) AND ((rat) OR (rats) OR (mouse) OR (mice))*.*”* Following this, the reference lists were screened for additional references that fit our analysis criteria. A filter was applied to the search so that only papers from the year 2000, in English, and excluding preprints were shown. Duplicate articles were identified and then removed. The articles returned in the search result were then individually evaluated if they were suitable to be included. Studies were excluded if they (i) reported no behavior data (the paper includes spinal cord injury but does not include any behavior data); (ii) no full-text available; (iii) lack of motor behavior data (the paper includes behavior data but not one of the accepted behavior measures); (iv) not a spinal cord injury model; (v) rehabilitation was not used; (vi) a lack of usable data (the study includes behavior data but data points are not able to be extracted e.g., from boxplots); (vii) a lack of an appropriate control group (i.e. a rehabilitation-only group receiving no additional experimental therapy); (viii) review articles; (ix) human subjects; (x) non-rodent subjects; (xi) juvenile animals used; (xii) brain or spinal cord stimulation.

### Extraction of graphical data

Following a full-text review, data extraction was performed and independently verified by three authors (J.M.G., R.M., and P.P.). The following information was extracted from each included study: name of the first author, year of publication, species and strain, type and subclassification of rehabilitation, injury type, level of injury, severity grade (as defined below), onset of rehabilitation, onset phase, duration per day of rehabilitation and total duration of the rehabilitation, sample numbers of the experimental and control groups, mean and standard deviation of behavior measures of the experimental and control groups. Only behavioral outcomes based on established protocols with demonstrated reproducibility across laboratories were extracted. This criterion led to the inclusion of the behavioral measures listed in **Table S1**. For graphical data, WebPlotDigitizer 5.0 was used for graphical data extraction. When mean and variability measures were only available in figures, the mean and either standard deviation (SD) or standard error of the mean (SEM) were extracted (**Fig. 7A**). When SEM was reported, the SD was calculated based on the corresponding group sample size (*n*) using the equation:

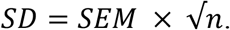

#### Classifications of graded study parameters

The severity grades were assigned using prespecified, study-specific classification criteria based on the extent of SCI and associated functional impairment. Classification integrated injury mechanism with locomotor recovery assessed using the Basso–Beattie–Bresnahan (BBB) locomotor rating scale (*88*).

- Grade 1 (mild): BBB score > 16 with only minor deficits in hindlimb plantar function. Typically observed following partial lesions (e.g., quadrant lesions) or very mild contusion injuries.
- Grade 2 (moderate): BBB score 12–16, with recovery of plantar placement. Animals generally regain weight-supported stepping but may show impaired coordination.
- Grade 3 (severe): High-force contusion injuries (typically ≥ 200 kdyn), characterized by failure to regain plantar placement in the affected hindlimbs.
- Grade 4 (most severe): Complete transection injuries, in which the spinal cord is fully severed and results in prolonged complete paralysis.

### Statistical analysis

#### Descriptive statistics and outlier removal

All analyses within this study were conducted using JASP (Version 0.98.1), running the *metafor* package in R. We systematically assessed potential outliers and influential observations before modelling. For the cervical and thoracic datasets, descriptive statistics, and visual diagnostics, including funnel plots, were inspected at the study level. Screening focused on continuous variables (e.g., SMDs, standard errors, sampling variances, and continuous moderators) and on categorical variables (e.g., moderator levels). For each continuous variable, we computed the mean, median, SD, minimum, maximum, and interquartile range (IQR). For categorical variables, frequency tables were generated to identify rare or unexpected categories and verify coding consistency. Missingness patterns were additionally examined to ensure that our findings were not driven by incomplete data.

We plotted histograms and boxplots for each continuous variable to visually assess distributional characteristics and identify potential extreme values. For key variable pairs (e.g., SMD vs. its standard error; SMD vs. continuous moderators), we used scatterplots to identify potential bivariate anomalies. These plots were inspected both overall and, where applicable, stratified by study or experimental subgroup to detect cluster-specific irregularities. For each continuous variable *x*, we computed standardized scores,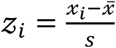, where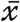 and *s* denote the sample mean and SD, respectively. Observations with *z*_*i*_ > ± 2.5were flagged as potential extreme values for further inspection. For variables showing substantial skewness, z-score flags were interpreted alongside distributional plots to avoid incorrectly classifying legitimate high- or low-value observations as outliers (**Figs. S1-2**).

#### Effect size calculation and modelling

For each comparison, the observed effect size was expressed as an SMD, *yi* with corresponding sampling variance *V*_*i*_. Meta-analyses were performed using multilevel random-effects meta-regression models fitted by restricted maximum likelihood (REML) using the rma.mv() function from the *metafor* package in R (**Fig. 2A,B**). REML was used to estimate variance components, as it provides approximately asymptotically unbiased estimates of variance components under random-effects assumptions and is widely recommended for multilevel meta-analysis (*29*). We report model coefficients (β), corresponding Wald-type z statistics, 95% confidence intervals, and associated *P* values. Model coefficients were evaluated using Wald-type z-tests with two-sided α = 0.05, and 95% confidence intervals were calculated accordingly `(test=“z”, level=95)`. We report model coefficients (β), corresponding z statistics and *P* values, the omnibus test of moderators (*Q*_*M*_), and the test of residual heterogeneity (*Q*_*E*_). Where appropriate, the proportion of the heterogeneity explained by moderators was quantified by comparing variance components between models.

#### Random-effects structure

To account for non-independence arising from multiple outcomes extracted from the same study, random intercepts were included at the study and experiment (`∼ 1 ∣Study/Experiment`). This hierarchical structure partitions residual heterogeneity into between-study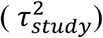 and between-experiment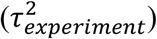 components while accounting for known sampling variance of each effect size. The model was expressed as:

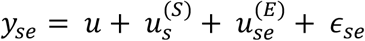

where:

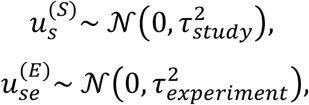

and:

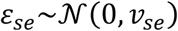

represents the sampling error associated with each effect size.

For moderator analyses, additional crossed random effects were included, where appropriate, to account for systematic variability associated with behavioral outcome measures and rehabilitation modalities. The model can be expressed as:

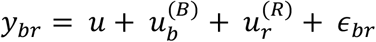

where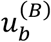 represents the random effect associated with behavioral outcome measure *b*,

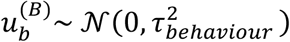

and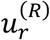 represents the random effect associated with rehabilitation modality *r*,

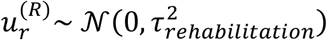

Here,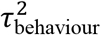 and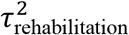denote the between-category variance components attributable to behavioral outcome measures and rehabilitation modalities, respectively, and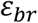 represents the residual error term

#### Meta-regressions

Potential sources of heterogeneity were investigated using mixed-effects meta-regression models fitted using REML in the *metafor* package. Moderators included rehabilitation type, rehabilitation onset phase, total rehabilitation duration (weeks), and daily rehabilitation exposure (minutes/day, **Fig. 2C**). The core random-effects structure accounted for dependence among effect sizes extracted from the same study and experiment using random intercepts for study and experiment levels.

For categorical moderators, effect sizes were modeled as:

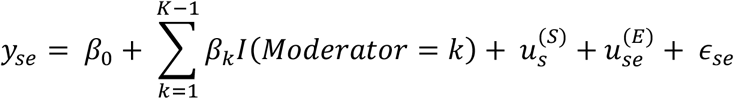

Where *I*(·) is an indicator variable identifying membership of a given category. In intercept models, β_0_ represents the estimated effect size of the reference category and β_k_ represents the difference in SMD units relative to the reference group. For selected analyses, no-intercept models were also fitted (−1 in the model formula), in which case each coefficient represented the estimated pooled effect size for the corresponding category.

For continuous moderators, the model was specified as:

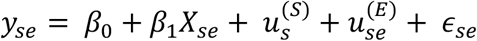

where *X*_s*e*_ denotes the moderator value for a given effect size (e.g., rehabilitation duration in weeks or minutes per day, **Fig. 2D**). The slope coefficient (*β*_1_) represents the expected change in SMD associated with a one-unit increase in the moderator (**Fig. 2E**).

To examine potential non-linear relationships between rehabilitation duration and treatment effect, quadratic meta-regression models were fitted:

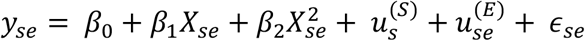

where *β*_2_ quantifies curvature in the duration-response relationship. A negative quadratic coefficient was interpreted as being consistent with diminishing returns or a plateau in treatment effect at longer intervention durations.

Moderator significance was assessed using omnibus Wald-type tests (*Q*_*M*_). Individual regression coefficients are reported as β estimates with corresponding standard errors, 95% confidence intervals, z statistics, and two-sided P values. Residual heterogeneity was assessed using the *Q*_*E*_ statistic. Regression coefficients are expressed in SMD units, with positive values indicating improved functional outcomes in rehabilitation-treated animals relative to controls.

#### Diagnostics

Where appropriate, funnel plots were visually inspected to evaluate potential small-study effects and publication bias. Given the multilevel structure of the dataset, with multiple effect sizes nested within experiments and studies, funnel plots were interpreted cautiously, as it may reflect true heterogeneity, dependence among effect sizes, or other study-level differences rather than publication bias alone. Therefore, funnel plots were considered qualitative diagnostic tools, and residual heterogeneity was assessed using the *Q*_*E*_ statistic from the multilevel meta-analytic models.

## List of Supplementary Materials

Supplementary Methods

Tables S1 to S12

Fig. S1 to Fig. S5

Supplementary References

Data file S1, which includes all characteristics, demographics, effect-size calculations, and citations of the studies included in the meta-analysis

## Acknowledgments

We would like to thank members of the Bradke and Griffin Labs for their ongoing support that led to this publication. We would specifically like to thank Norbert Weidner, Sina Stern, Botao Tan, and Andreas Husch for critically reviewing our manuscript.

## Funding

This work was supported in part by:

The CatWalk Spinal Cord Injury Research Trust: Postdoctoral Research Fellow (JMG) Wings for Life project 300 (JMG, FB)

The International Foundation for Research in Paraplegia (FB),

German Research Foundation SFB1089:227953431(JMG, FB); SFB1158 (FB); SFB1451 (FB); SFB1690 (FB); SPP2395 (FB); RTG1960 (FB); and C3NS (FB)

ERANET: Axon repair and Rater SCI (FB) ibehave Network Nordrhein-Westfalen (FB) European Research Council # 101041677 (AA).

## Author contributions

Conceptualization: JMG

Methodology: JMG, AA, CJ

Article screening: JMG, RM, PP

Data extraction: JMG, RM, PP

Analysis: JMG, AA

Supervision: FB

Writing – original draft: JMG

Writing – review & editing: JMG, AA, PP, SK, OC, DS, CJ, FB

## Data and materials availability

All data associated with this study are present in the paper or Supplementary Materials. The data used to perform the analyses and the citations for all studies included in the meta-analysis are available in supplementary data file S1.

