## Supplementary Materials for "Voluntary engagement predicts recovery across rehabilitation time points after experimental spinal cord injury"

Jarred M. Griffin *et al.*

**The PDF file includes:**

Supplementary Methods
Tables S1 to S12
Figs. S1 to S5
Supplementary References

#### Supplementary Methods:

##### Classification of stress grades

Rehabilitation interventions were classified according to their level of perceived stress burden using a four-point ordinal scale. Grade 0 comprised passive or voluntary activity-based approaches, including environmental enrichment and voluntary wheel running. Grade 1 included non-weight-supported quadrupedal training modalities such as treadmill training, forced wheel running, staircase climbing, exercise ball training, runway tasks, maze navigation, and ramp walking. Grade 2 comprised more intensive locomotor rehabilitation approaches involving weight-supported and non-weight-supported bipedal treadmill training, weight-supported quadrupedal treadmill training, and swimming. Grade 3 represented the most complex rehabilitation interventions and included cycling, water stepping, rotarod training, ladder and grid walking tasks, and combined swimming plus exercise ball training. Higher grades therefore reflected increasing levels of motor challenge, task specificity, and rehabilitation intensity.

Stress coefficient equation:

$$sc = \frac{sg \times t}{10}$$

Where ‘sc’ is the stress coefficient, ‘sg’ is the stress grade and ‘t’ is the daily duration spent engaged in the rehabilitation task.

#### Supplementary Tables

**Table S1. List of accepted behavioral measures used in the meta-analysis and their description.** We identified 12 behavioral measures that would be acceptable for inclusion within this meta-analysis. These are based on comparable and standardized behavior measures used in the field of spinal cord injury. A description of each measure is presented, alongside a citation for each one.

| Behavior Measure | Description | Example citation |
|---|---|---|
| **Basso-Beattie-Bresnahan scale (BBB)** | Observational open field test based on grading hindlimb locomotion of rats from 0 (no spontaneous activity) to 21 (normal movement). The BBB scale is the standard behavioral measure for the field of SCI research | (*1*) |
| **Basso Mouse Scale (BMS)** | Observational open field test based on grading hindlimb locomotion of mice from 0 (no spontaneous activity) to 9 (normal movement). | (*2*) |
| **Pellet reaching success** | Assesses skilled forepaw use and motor functioning in the ability of a rat to grasp a sugar pellet successfully. Included in this category are the window reaching and staircase reaching. | (*3*) |
| **Forepaw and hindpaw ladder/grid score** | Assesses fore- and hindlimb stepping, placing and coordination. Included in this is the standard horizontal error ladder as well as | (*4*) |

| | | |
|---|---|---|
| | the grid test. Accepted measure is the percentage slips of total steps | |
| **Swim score** | Louisville swim scale. Evaluates three characteristics of swimming that are highly altered by spinal cord injury—namely, hindlimb movement, forelimb dependency, and body position | (*5*) |
| **Narrow beam score** | The method used to assess the ability of rats to balance on 1-m-long elevated beams with different cross-sectional profiles. | (*6*) |
| **Coordination** | Gait analysis system for assessing subtle differences in gait; animals are filmed from below while crossing a glass walkway, and paw print locations are recorded and subsequently analyzed with the Catwalk software. | (*7*) |
| **Paw rotation** | The measurement of the degree of paw rotation from the parallel to the body line. The measurement can be made from the CatWalk Gait XT analysis system or manually from paw print makers. | (*7*) |
| **Rotarod score** | Measure of locomotor performance, particularly motor coordination, in which mice and rats are placed on an elevated revolving rod and the time it takes them to fall is recorded. | (*8*) |
| **Stride length** | The absolute measurement of the stride length, typically calculated from the CatWalk Gait XT gait analysis system can be determined by other catwalk systems or manuals from print markings. | (*7*) |
| **Average Combined Score (ACOS)** | Semi-quantitative method to assess hindlimb movement in mice; combines non-locomotor movement, locomotor-like movement, and movement amplitude, to quantify and score hindlimb movement. | (*9*) |
| **Tarlov scale** | Observational 4-point scales similar to the ASIA impairment scale used to evaluate the movement, weight-bearing, and walking abilities of the hind limbs | (*10*) |

**Table S2. Model summary from multilevel meta-analysis of the effects of rehabilitation following cervical spinal cord injury stratified by behavior measures.**

#### Model Summary

*Meta-Analytic Tests*

| | Subgroup | Test | *P* |
|---|---|---|---|
| Heterogeneity | Forepaw ladder/grid | $Q_e(27) = 57.22$ | < .001 |
| | Reaching success | $Q_e(19) = 38.10$ | .006 |
| | BBB/BMS | $Q_e(6) = 5.26$ | .511 |
| | Hindpaw ladder/grid | $Q_e(13) = 10.03$ | .691 |
| | Narrow beam | | |
| | Stride length | | |
| | Coordination | | |
| | Rotarod | | |
| Pooled effect | Forepaw ladder/grid | z = 3.60 | < .001 |
| | Reaching success | z = 3.57 | < .001 |
| | BBB/BMS | z = 2.10 | .035 |
| | Hindpaw ladder/grid | z = 3.25 | .001 |
| Subgroup differences | | $Q_m(3) = 1.37$ | .713 |

*Meta-Analytic Estimates*

| | Subgroup | Estimate | Standard Error | 95% CI Lower | 95% CI Upper | 95% PI Lower | 95% PI Upper |
|---|---|---|---|---|---|---|---|
| Pooled effect | Forepaw ladder/grid | 0.5093 | 0.1415 | 0.2321 | 0.7866 | -0.4889 | 1.508 |
| | Reaching success | 0.5675 | 0.1589 | 0.2561 | 0.8790 | -0.4278 | 1.563 |
| | BBB/BMS | 0.3479 | 0.1653 | 0.02392 | 0.6719 | -0.1157 | 0.8115 |
| | Hindpaw ladder/grid | 0.3883 | 0.1196 | 0.1539 | 0.6227 | 0.1539 | 0.6227 |

**Table S3. Model summary from multilevel meta-analysis of the effects of rehabilitation following thoracic spinal cord injury stratified by behavior measures.**

#### Model Summary

*Meta-Analytic Tests*

| | Subgroup | Test | *P* |
|---|---|---|---|
| Heterogeneity | BBB/BMS | $Q_e(135) = 360.90$ | < .001 |
| | Narrow beam | $Q_e(1) = 0.04$ | .846 |
| | Hindpaw ladder/grid | $Q_e(35) = 73.86$ | < .001 |
| | Coordination | $Q_e(12) = 44.98$ | < .001 |
| | Swim score | $Q_e(3) = 6.93$ | .074 |
| | Paw rotation | $Q_e(6) = 8.03$ | .236 |
| | Stride length | $Q_e(10) = 52.07$ | < .001 |
| | Forepaw ladder/grid | $Q_e(1) = 0.79$ | .373 |
| | ACOS/Tarlov | $Q_e(2) = 0.05$ | .976 |
| | Rotarod | $Q_e(1) = 0.20$ | .653 |
| Pooled effect | BBB/BMS | z = 10.36 | < .001 |
| | Narrow beam | z = -0.31 | .757 |
| | Hindpaw ladder/grid | z = 6.87 | < .001 |
| | Coordination | z = 2.06 | .040 |
| | Swim score | z = 2.12 | .034 |
| | Paw rotation | z = 5.39 | < .001 |
| | Stride length | z = 2.24 | .025 |
| | Forepaw ladder/grid | z = 3.92 | < .001 |
| | ACOS/Tarlov | z = 4.03 | < .001 |
| | Rotarod | z = 3.88 | < .001 |
| Subgroup differences | | $Q_m(9) = 15.31$ | .083 |

*Meta-Analytic Estimates*

| | Subgroup | Estimate | Standard Error | 95% CI Lower | 95% CI Upper | 95% PI Lower | 95% PI Upper |
|---|---|---|---|---|---|---|---|
| Pooled effect | BBB/BMS | 0.8261 | 0.07976 | 0.6698 | 0.9824 | -0.5180 | 2.170 |
| | Narrow beam | -0.1213 | 0.3927 | -0.8910 | 0.6484 | -0.8910 | 0.6484 |

*Meta-Analytic Estimates*

| Subgroup | Estimate | Standard Error | 95% CI Lower | 95% CI Upper | 95% PI Lower | 95% PI Upper |
|---|---|---|---|---|---|---|
| Hindpaw ladder/grid | 0.9021 | 0.1312 | 0.6449 | 1.159 | -0.2033 | 2.008 |
| Coordination | 0.6186 | 0.3009 | 0.02884 | 1.208 | -1.088 | 2.325 |
| Swim score | 1.014 | 0.4787 | 0.07554 | 1.952 | -0.6736 | 2.701 |
| Paw rotation | 1.104 | 0.2047 | 0.7029 | 1.505 | 0.7029 | 1.505 |
| Stride length | 0.8844 | 0.3951 | 0.1100 | 1.659 | -1.468 | 3.237 |
| Forepaw ladder/grid | 1.345 | 0.3433 | 0.6717 | 2.017 | 0.6717 | 2.017 |
| ACOS/Tarlov | 1.276 | 0.3168 | 0.6548 | 1.896 | 0.6548 | 1.896 |
| Rotarod | 1.570 | 0.4050 | 0.7765 | 2.364 | 0.7765 | 2.364 |

**Table S4. Meta-Regression Summary for the cervical data stratified by behavior measure with rehabilitation type as a categorical moderator**

#### Model Summary

*Meta-Analytic Tests*

| | Subgroup | Test | *P* |
|---|---|---|---|
| Residual heterogeneity | Forepaw ladder/grid | $Q_e(21) = 34.47$ | .032 |
| | Reaching success | $Q_e(17) = 27.20$ | .055 |
| | BBB/BMS | $Q_e(3) = 1.14$ | .768 |
| | Hindpaw ladder/grid | $Q_e(9) = 5.08$ | .827 |
| Pooled effect | Forepaw ladder/grid | z = 3.91 | < .001 |
| | Reaching success | z = 4.16 | < .001 |
| | BBB/BMS | z = 2.80 | .005 |
| | Hindpaw ladder/grid | z = 3.17 | .002 |
| Moderation | Forepaw ladder/grid | $Q_m(7) = 26.71$ | < .001 |
| | Reaching success | $Q_m(3) = 24.63$ | < .001 |
| | BBB/BMS | $Q_m(4) = 9.07$ | .059 |
| | Hindpaw ladder/grid | $Q_m(5) = 15.50$ | .008 |
| Subgroup differences | | $Q_m(3) = 1.09$ | .780 |

*Meta-Analytic Estimates*

| | Subgroup | Estimate | Standard Error | 95% CI Lower | 95% CI Upper | 95% PI Lower | 95% PI Upper |
|---|---|---|---|---|---|---|---|
| Pooled effect | Forepaw ladder/grid | 0.5029 | 0.1285 | 0.2510 | 0.7547 | -0.3281 | 1.334 |
| | Reaching success | 0.5616 | 0.1351 | 0.2968 | 0.8264 | -0.1704 | 1.294 |
| | BBB/BMS | 0.4358 | 0.1556 | 0.1308 | 0.7407 | 0.1308 | 0.7407 |
| | Hindpaw ladder/grid | 0.3834 | 0.1209 | 0.1464 | 0.6203 | 0.1464 | 0.6203 |

*Effect Size Meta-Regression Coefficients*

| | Subgroup | Estimate | Standard Error | 95% CI Lower | 95% CI Upper | z | *P* |
|---|---|---|---|---|---|---|---|
| Rehabilitation type (Environmental enrichment) | Forepaw ladder/grid | 0.9708 | 0.3150 | 0.3533 | 1.588 | 3.082 | .002 |
| | Reaching success | -0.4794 | 0.4136 | -1.290 | 0.3313 | -1.159 | .246 |
| | BBB/BMS | 0.5248 | 0.4152 | -0.2890 | 1.339 | 1.264 | .206 |
| | Hindpaw ladder/grid | 0.6274 | 0.3515 | -0.06150 | 1.316 | 1.785 | .074 |
| Rehabilitation type (Ladder/grid) | Forepaw ladder/grid | 0.9913 | 0.3905 | 0.2258 | 1.757 | 2.538 | .011 |
| | BBB/BMS | -0.08124 | 0.2541 | -0.5793 | 0.4168 | -0.3197 | .749 |
| Rehabilitation type (Overground sensorimotor) | Forepaw ladder/grid | 1.432 | 0.6438 | 0.1698 | 2.693 | 2.224 | .026 |
| | BBB/BMS | 0.7774 | 0.4638 | -0.1316 | 1.686 | 1.676 | .094 |
| Rehabilitation type (Quadrupedal treadmill) | Forepaw ladder/grid | 0.5242 | 0.2929 | -0.04987 | 1.098 | 1.790 | .074 |
| | Hindpaw ladder/grid | 0.5031 | 0.1731 | 0.1639 | 0.8423 | 2.907 | .004 |
| Rehabilitation type (Quadrupedal treadmill and reach training) | Forepaw ladder/grid | 0.8243 | 0.6591 | -0.4675 | 2.116 | 1.251 | .211 |
| | Reaching success | 1.352 | 0.6545 | 0.06900 | 2.634 | 2.065 | .039 |
| | Hindpaw ladder/grid | -0.4878 | 0.5074 | -1.482 | 0.5066 | -0.9614 | .336 |
| Rehabilitation type (Reach training) | Forepaw ladder/grid | 0.2567 | 0.1808 | -0.09771 | 0.6112 | 1.420 | .156 |
| | Reaching success | 0.6376 | 0.1463 | 0.3509 | 0.9242 | 4.359 | < .001 |
| | Hindpaw ladder/grid | 0.2133 | 0.2192 | -0.2163 | 0.6429 | 0.9732 | .330 |
| Rehabilitation type (Running wheel) | Forepaw ladder/grid | -0.5245 | 0.6773 | -1.852 | 0.8030 | -0.7744 | .439 |
| | BBB/BMS | 0.4573 | 0.2142 | 0.03743 | 0.8772 | 2.135 | .033 |
| | Hindpaw ladder/grid | 0.7282 | 0.5163 | -0.2838 | 1.740 | 1.410 | .158 |

**Table S5. Meta-Regression Summary for the thoracic data subgrouped by rehabilitation type**

#### Model Summary

*Meta-Analytic Tests*

| | Subgroup | Test | *P* |
|---|---|---|---|
| Residual heterogeneity | BBB/BMS | $Q_e(122) = 296.60$ | < .001 |
| | Narrow beam | $Q_e(0) = 0.00$ | 1.000 |
| | Hindpaw ladder/grid | $Q_e(26) = 60.37$ | < .001 |
| | Coordination | $Q_e(6) = 39.49$ | < .001 |
| | Swim score | $Q_e(1) = 2.66$ | .103 |
| | Paw rotation | $Q_e(3) = 0.14$ | .987 |
| | Stride length | $Q_e(6) = 36.42$ | < .001 |
| | Forepaw ladder/grid | $Q_e(0) = 0.00$ | 1.000 |
| | ACOS/Tarlov | $Q_e(0) = 0.00$ | 1.000 |
| | Rotarod | $Q_e(0) = 0.00$ | 1.000 |
| Pooled effect | BBB/BMS | z = 10.89 | < .001 |
| | Narrow beam | z = -0.29 | .769 |
| | Hindpaw ladder/grid | z = 6.57 | < .001 |
| | Coordination | z = 1.59 | .112 |
| | Swim score | z = 2.24 | .025 |
| | Paw rotation | z = 6.01 | < .001 |
| | Stride length | z = 2.10 | .036 |
| | Forepaw ladder/grid | z = 3.82 | < .001 |
| | ACOS/Tarlov | z = 4.03 | < .001 |
| | Rotarod | z = 3.90 | < .001 |
| Moderation | BBB/BMS | $Q_m(14) = 145.62$ | < .001 |
| | Narrow beam | $Q_m(2) = 0.13$ | .936 |
| | Hindpaw ladder/grid | $Q_m(10) = 47.87$ | < .001 |
| | Coordination | $Q_m(7) = 4.95$ | .666 |
| | Swim score | $Q_m(3) = 5.81$ | .121 |
| | Paw rotation | $Q_m(4) = 36.99$ | < .001 |
| | Stride length | $Q_m(5) = 7.08$ | .215 |
| | Forepaw ladder/grid | $Q_m(2) = 16.13$ | < .001 |
| | ACOS/Tarlov | $Q_m(3) = 16.27$ | .001 |
| | Rotarod | $Q_m(2) = 15.23$ | < .001 |
| Subgroup differences | | $Q_m(9) = 19.17$ | .024 |

*Meta-Analytic Estimates*

| | | | | 95% CI | | 95% PI | |
|---|---|---|---|---|---|---|---|
| | Subgroup | Estimate | Standard Error | Lower | Upper | Lower | Upper |
| Pooled effect | BBB/BMS | 0.8138 | 0.07474 | 0.6673 | 0.9603 | -0.4285 | 2.056 |
| | Narrow beam | -0.1156 | 0.3938 | -0.8875 | 0.6563 | -0.8875 | 0.6563 |
| | Hindpaw ladder/grid | 0.9113 | 0.1386 | 0.6396 | 1.183 | -0.3073 | 2.130 |
| | Coordination | 0.6328 | 0.3979 | -0.1470 | 1.413 | -1.718 | 2.983 |
| | Swim score | 1.182 | 0.5284 | 0.1467 | 2.218 | -0.7137 | 3.079 |
| | Paw rotation | 1.329 | 0.2212 | 0.8954 | 1.763 | 0.8954 | 1.763 |
| | Stride length | 0.8766 | 0.4174 | 0.05842 | 1.695 | -1.637 | 3.390 |

*Meta-Analytic Estimates*

| | | | 95% CI | | 95% PI | |
|---|---|---|---|---|---|---|
| Subgroup | Estimate | Standard Error | Lower | Upper | Lower | Upper |
| Forepaw ladder/grid | 1.526 | 0.3991 | 0.7437 | 2.308 | 0.7437 | 2.308 |
| ACOS/Tarlov | 1.277 | 0.3168 | 0.6560 | 1.898 | 0.6560 | 1.898 |
| Rotarod | 1.580 | 0.4056 | 0.7852 | 2.375 | 0.7852 | 2.375 |

*Effect Size Meta-Regression Coefficients*

| | | | | 95% CI | | | |
|---|---|---|---|---|---|---|---|
| | Subgroup | Estimate | Standard Error | Lower | Upper | z | *P* |
| Rehabilitation type (Bipedal treadmill) | BBB/BMS | 0.6008 | 0.1571 | 0.2930 | 0.9087 | 3.825 | < .001 |
| | Hindpaw ladder/grid | 0.8898 | 0.4593 | -0.01044 | 1.790 | 1.937 | .053 |
| | Coordination | 1.021 | 0.6012 | -0.1571 | 2.200 | 1.699 | .089 |
| | Paw rotation | 0.7687 | 0.3555 | 0.07193 | 1.465 | 2.162 | .031 |
| | Stride length | 0.8717 | 0.5572 | -0.2203 | 1.964 | 1.565 | .118 |
| | ACOS/Tarlov | 1.367 | 0.5553 | 0.2783 | 2.455 | 2.461 | .014 |
| Rehabilitation type (Cycling) | BBB/BMS | 0.01268 | 0.3044 | -0.5839 | 0.6092 | 0.04165 | .967 |
| | Hindpaw ladder/grid | 0.3925 | 0.8409 | -1.256 | 2.041 | 0.4667 | .641 |
| | Paw rotation | 1.185 | 0.4849 | 0.2347 | 2.135 | 2.444 | .015 |
| Rehabilitation type (Environmental enrichment) | BBB/BMS | 1.247 | 0.2074 | 0.8403 | 1.653 | 6.012 | < .001 |
| | Hindpaw ladder/grid | 1.218 | 0.2555 | 0.7177 | 1.719 | 4.769 | < .001 |
| | Coordination | 0.6552 | 0.8605 | -1.031 | 2.342 | 0.7614 | .446 |
| | Swim score | 0.3512 | 0.9173 | -1.447 | 2.149 | 0.3829 | .702 |
| | Paw rotation | 2.467 | 0.5420 | 1.405 | 3.529 | 4.552 | < .001 |
| | Stride length | 1.404 | 1.373 | -1.287 | 4.095 | 1.023 | .307 |
| | Forepaw ladder/grid | 1.881 | 0.6934 | 0.5224 | 3.240 | 2.713 | .007 |
| Rehabilitation type (Exercise ball) | BBB/BMS | 1.006 | 0.7206 | -0.4058 | 2.419 | 1.397 | .162 |
| Rehabilitation type (Ladder/grid) | BBB/BMS | 0.3614 | 0.8064 | -1.219 | 1.942 | 0.4482 | .654 |
| | Hindpaw ladder/grid | -0.1497 | 0.7862 | -1.691 | 1.391 | -0.1904 | .849 |
| Rehabilitation type (Maze) | BBB/BMS | 0.3148 | 0.7547 | -1.164 | 1.794 | 0.4172 | .677 |
| Rehabilitation type (Quadrupedal treadmill) | BBB/BMS | 0.9524 | 0.1190 | 0.7192 | 1.186 | 8.004 | < .001 |
| | Narrow beam | -0.1922 | 0.5358 | -1.242 | 0.8579 | -0.3587 | .720 |

*Effect Size Meta-Regression Coefficients*

| | Subgroup | Estimate | Standard Error | 95% CI Lower | 95% CI Upper | z | *P* |
|---|---|---|---|---|---|---|---|
| | Hindpaw ladder/grid | 0.8363 | 0.2379 | 0.3700 | 1.303 | 3.515 | < .001 |
| | Coordination | 0.3503 | 0.7504 | -1.121 | 1.821 | 0.4668 | .641 |
| | Paw rotation | 0.8232 | 0.3473 | 0.1425 | 1.504 | 2.370 | .018 |
| | Stride length | 0.8218 | 0.7044 | -0.5588 | 2.202 | 1.167 | .243 |
| | Forepaw ladder/grid | 1.170 | 0.3952 | 0.3957 | 1.945 | 2.961 | .003 |
| | ACOS/Tarlov | 1.268 | 0.5480 | 0.1942 | 2.342 | 2.314 | .021 |
| | Rotarod | 1.763 | 0.5891 | 0.6079 | 2.917 | 2.992 | .003 |
| Rehabilitation type (Ramp) | BBB/BMS | 1.122 | 0.7874 | -0.4212 | 2.665 | 1.425 | .154 |
| Rehabilitation type (Rotarod Training) | BBB/BMS | -1.555 | 0.9112 | -3.341 | 0.2308 | -1.707 | .088 |
| | Coordination | 0.4248 | 1.221 | -1.968 | 2.818 | 0.3479 | .728 |
| Rehabilitation type (Running wheel) | BBB/BMS | 1.115 | 0.2645 | 0.5965 | 1.633 | 4.215 | < .001 |
| | Hindpaw ladder/grid | 0.7135 | 0.3424 | 0.04245 | 1.385 | 2.084 | .037 |
| | Coordination | 1.010 | 0.9637 | -0.8790 | 2.899 | 1.048 | .295 |
| | Rotarod | 1.398 | 0.5577 | 0.3048 | 2.491 | 2.506 | .012 |
| Rehabilitation type (Runway) | BBB/BMS | 0.4063 | 0.5603 | -0.6920 | 1.504 | 0.7250 | .468 |
| | Hindpaw ladder/grid | 0.5221 | 0.7911 | -1.028 | 2.073 | 0.6600 | .509 |
| Rehabilitation type (Staircase climbing) | BBB/BMS | 0.000 | 0.7602 | -1.490 | 1.490 | 0.000 | 1.000 |
| | Hindpaw ladder/grid | 1.636 | 0.7810 | 0.1048 | 3.166 | 2.094 | .036 |
| | Coordination | 0.8759 | 1.216 | -1.508 | 3.260 | 0.7202 | .471 |
| | Stride length | 2.182 | 1.327 | -0.4180 | 4.782 | 1.645 | .100 |
| Rehabilitation type (Swimming) | BBB/BMS | 0.9275 | 0.2790 | 0.3808 | 1.474 | 3.325 | < .001 |
| | Hindpaw ladder/grid | 0.9020 | 0.7666 | -0.6005 | 2.405 | 1.177 | .239 |
| | Swim score | 0.9832 | 0.7256 | -0.4390 | 2.405 | 1.355 | .175 |
| | ACOS/Tarlov | 1.196 | 0.5428 | 0.1318 | 2.260 | 2.203 | .028 |
| Rehabilitation type (Water stepping) | BBB/BMS | 0.4984 | 0.3423 | -0.1725 | 1.169 | 1.456 | .145 |
| Rehabilitation type (Swimming + exercise ball) | Narrow beam | -0.03899 | 0.5774 | -1.171 | 1.093 | -0.06753 | .946 |
| | Hindpaw ladder/grid | 0.6663 | 0.8481 | -0.9958 | 2.328 | 0.7857 | .432 |
| | Coordination | -0.6044 | 1.276 | -3.106 | 1.897 | -0.4737 | .636 |
| | Stride length | -0.7677 | 1.352 | -3.418 | 1.882 | -0.5678 | .570 |

*Effect Size Meta-Regression Coefficients*

| | Subgroup | Estimate | Standard Error | 95% CI Lower | 95% CI Upper | z | *P* |
|---|---|---|---|---|---|---|---|
| Rehabilitation type (Environmental enrichment + swimming) | Swim score | 2.412 | 1.233 | -0.004549 | 4.829 | 1.956 | .050 |

#### Table S6. Meta-Regression Summary for the thoracic data subgrouped by rehabilitation type including subcategories

#### Model Summary

*Meta-Analytic Tests*

| | Subgroup | Test | *P* |
|---|---|---|---|
| Residual heterogeneity | BBB/BMS | $Q_e(113) = 267.22$ | < .001 |
| | Narrow beam | $Q_e(0) = 0.00$ | 1.000 |
| | Hindpaw ladder/grid | $Q_e(23) = 56.63$ | < .001 |
| | Coordination | $Q_e(4) = 20.24$ | < .001 |
| | Swim score | $Q_e(1) = 2.66$ | .103 |
| | Paw rotation | $Q_e(2) = 0.07$ | .967 |
| | Stride length | $Q_e(4) = 23.43$ | < .001 |
| | Forepaw ladder/grid | $Q_e(0) = 0.00$ | 1.000 |
| | ACOS/Tarlov | $Q_e(0) = 0.00$ | 1.000 |
| | Rotarod | $Q_e(0) = 0.00$ | 1.000 |
| Pooled effect | BBB/BMS | z = 10.83 | < .001 |
| | Narrow beam | z = -0.29 | .769 |
| | Hindpaw ladder/grid | z = 6.39 | < .001 |
| | Coordination | z = 1.51 | .132 |
| | Swim score | z = 2.24 | .025 |
| | Paw rotation | z = 5.99 | < .001 |
| | Stride length | z = 1.96 | .050 |
| | Forepaw ladder/grid | z = 3.82 | < .001 |
| | ACOS/Tarlov | z = 4.03 | < .001 |
| | Rotarod | z = 3.90 | < .001 |
| Moderation | BBB/BMS | $Q_m(20) = 151.65$ | < .001 |
| | Narrow beam | $Q_m(2) = 0.13$ | .936 |
| | Hindpaw ladder/grid | $Q_m(13) = 46.43$ | < .001 |
| | Coordination | $Q_m(9) = 14.38$ | .109 |
| | Swim score | $Q_m(3) = 5.81$ | .121 |
| | Paw rotation | $Q_m(5) = 37.06$ | < .001 |
| | Stride length | $Q_m(7) = 7.75$ | .355 |
| | Forepaw ladder/grid | $Q_m(2) = 16.13$ | < .001 |
| | ACOS/Tarlov | $Q_m(3) = 16.27$ | .001 |
| | Rotarod | $Q_m(2) = 15.23$ | < .001 |
| Subgroup differences | | $Q_m(9) = 19.82$ | .019 |

*Meta-Analytic Estimates*

| | Subgroup | Estimate | Standard Error | 95% CI Lower | 95% CI Upper | 95% PI Lower | 95% PI Upper |
|---|---|---|---|---|---|---|---|
| Pooled effect | BBB/BMS | 0.8010 | 0.07399 | 0.6560 | 0.9461 | -0.4104 | 2.012 |
| | Narrow beam | -0.1156 | 0.3938 | -0.8875 | 0.6563 | -0.8875 | 0.6563 |
| | Hindpaw ladder/grid | 0.9140 | 0.1430 | 0.6339 | 1.194 | -0.3752 | 2.203 |
| | Coordination | 0.5712 | 0.3795 | -0.1726 | 1.315 | -1.715 | 2.857 |
| | Swim score | 1.182 | 0.5284 | 0.1467 | 2.218 | -0.7137 | 3.079 |
| | Paw rotation | 1.327 | 0.2214 | 0.8925 | 1.760 | 0.8925 | 1.760 |
| | Stride length | 0.8331 | 0.4258 | -0.001419 | 1.668 | -1.809 | 3.476 |
| | Forepaw ladder/grid | 1.526 | 0.3991 | 0.7437 | 2.308 | 0.7437 | 2.308 |
| | ACOS/Tarlo v | 1.277 | 0.3168 | 0.6560 | 1.898 | 0.6560 | 1.898 |
| | Rotarod | 1.580 | 0.4056 | 0.7852 | 2.375 | 0.7852 | 2.375 |

*Effect Size Meta-Regression Coefficients*

| | Subgroup | Estimate | Standard Error | 95% CI Lower | 95% CI Upper | z | *P* |
|---|---|---|---|---|---|---|---|
| Subclassificati on✻ ) | BBB/BMS | 0.9030 | 0.3616 | 0.1943 | 1.612 | 2.497 | .013 |
| | Forepaw ladder/grid | 1.170 | 0.3952 | 0.3957 | 1.945 | 2.961 | .003 |
| Subclassificati on (Exercise ball) | BBB/BMS | 1.018 | 0.6978 | -0.3495 | 2.386 | 1.459 | .145 |
| Subclassificati on (Forced running wheel) | BBB/BMS | 1.737 | 0.5055 | 0.7465 | 2.728 | 3.437 | < .001 |
| Subclassificati on (General enrichment) | BBB/BMS | 0.8090 | 0.2748 | 0.2704 | 1.347 | 2.944 | .003 |
| | Hindpaw ladder/grid | 0.9319 | 0.3434 | 0.2588 | 1.605 | 2.714 | .007 |
| | Coordination | 0.6543 | 0.8417 | -0.9955 | 2.304 | 0.7773 | .437 |
| Subclassificati on (Horizontal ladder) | BBB/BMS | 0.3614 | 0.7941 | -1.195 | 1.918 | 0.4551 | .649 |
| | Hindpaw ladder/grid | -0.1497 | 0.8142 | -1.746 | 1.446 | -0.1838 | .854 |
| Subclassificati on (Maze) | BBB/BMS | 0.2942 | 0.7494 | -1.175 | 1.763 | 0.3926 | .695 |
| Subclassificati on (Non-weight supported bipedal treadmill) | BBB/BMS | 0.6147 | 0.4291 | -0.2262 | 1.456 | 1.433 | .152 |
| | Hindpaw ladder/grid | 0.7298 | 0.7681 | -0.7757 | 2.235 | 0.9501 | .342 |

*Effect Size Meta-Regression Coefficients*

| | Subgroup | Estimate | Standard Error | 95% CI Lower | 95% CI Upper | z | *P* |
|---|---|---|---|---|---|---|---|
| | Stride length | 2.042 | 1.368 | -0.6383 | 4.723 | 1.493 | .135 |
| Subclassification (Non-weight supported quadrupedal treadmill) | BBB/BMS | 0.9010 | 0.1483 | 0.6103 | 1.192 | 6.074 | < .001 |
| | Narrow beam | -0.1922 | 0.5358 | -1.242 | 0.8579 | -0.3587 | .720 |
| | Hindpaw ladder/grid | 0.8380 | 0.2901 | 0.2695 | 1.407 | 2.889 | .004 |
| | Coordination | -0.7021 | 1.197 | -3.048 | 1.644 | -0.5867 | .557 |
| | Stride length | 1.244 | 1.011 | -0.7384 | 3.226 | 1.230 | .219 |
| | ACOS/Tarlov | 1.268 | 0.5480 | 0.1942 | 2.342 | 2.314 | .021 |
| | Rotarod | 1.763 | 0.5891 | 0.6079 | 2.917 | 2.992 | .003 |
| Subclassification (Passive cycling) | BBB/BMS | -0.03220 | 0.3052 | -0.6303 | 0.5659 | -0.1055 | .916 |
| | Hindpaw ladder/grid | 0.3925 | 0.8672 | -1.307 | 2.092 | 0.4526 | .651 |
| | Paw rotation | 1.185 | 0.4849 | 0.2347 | 2.135 | 2.444 | .015 |
| Subclassification (Ramp) | BBB/BMS | 1.101 | 0.7823 | -0.4318 | 2.635 | 1.408 | .159 |
| Subclassification (Rotarod Training) | BBB/BMS | -1.555 | 0.9004 | -3.320 | 0.2097 | -1.727 | .084 |
| | Coordination | -0.7274 | 1.254 | -3.184 | 1.730 | -0.5802 | .562 |
| Subclassification (Runway) | BBB/BMS | 0.4057 | 0.5515 | -0.6752 | 1.487 | 0.7357 | .462 |
| | Hindpaw ladder/grid | 0.5221 | 0.8190 | -1.083 | 2.127 | 0.6375 | .524 |
| Subclassification (Staircase climbing) | BBB/BMS | 0.000 | 0.7473 | -1.465 | 1.465 | 0.000 | 1.000 |
| | Hindpaw ladder/grid | 1.636 | 0.8092 | 0.04947 | 3.222 | 2.021 | .043 |
| | Coordination | 0.8759 | 1.190 | -1.456 | 3.208 | 0.7363 | .462 |
| | Stride length | 2.182 | 1.388 | -0.5382 | 4.903 | 1.572 | .116 |
| Subclassification (Swimming) | BBB/BMS | 0.9862 | 0.2957 | 0.4066 | 1.566 | 3.335 | < .001 |
| | Hindpaw ladder/grid | 0.9020 | 0.7954 | -0.6569 | 2.461 | 1.134 | .257 |
| | Swim score | 0.9832 | 0.7256 | -0.4390 | 2.405 | 1.355 | .175 |
| | ACOS/Tarlov | 1.196 | 0.5428 | 0.1318 | 2.260 | 2.203 | .028 |
| Subclassification (Targeted enrichment) | BBB/BMS | 1.775 | 0.3077 | 1.172 | 2.378 | 5.769 | < .001 |
| | Hindpaw ladder/grid | 1.623 | 0.4119 | 0.8161 | 2.431 | 3.941 | < .001 |
| | Swim score | 0.3512 | 0.9173 | -1.447 | 2.149 | 0.3829 | .702 |
| | Paw rotation | 2.467 | 0.5420 | 1.405 | 3.529 | 4.552 | < .001 |
| | Stride length | 1.404 | 1.432 | -1.403 | 4.212 | 0.9802 | .327 |

*Effect Size Meta-Regression Coefficients*

| | Subgroup | Estimate | Standard Error | 95% CI Lower | 95% CI Upper | z | *P* |
|---|---|---|---|---|---|---|---|
| | Forepaw ladder/grid | 1.881 | 0.6934 | 0.5224 | 3.240 | 2.713 | .007 |
| Subclassification (Voluntary running wheel) | BBB/BMS | 0.8044 | 0.3046 | 0.2074 | 1.401 | 2.641 | .008 |
| | Hindpaw ladder/grid | 0.7088 | 0.3281 | 0.06575 | 1.352 | 2.160 | .031 |
| | Coordination | 0.6320 | 1.135 | -1.592 | 2.856 | 0.5571 | .577 |
| | Rotarod | 1.398 | 0.5577 | 0.3048 | 2.491 | 2.506 | .012 |
| Subclassification (Water stepping) | BBB/BMS | 0.4796 | 0.3292 | -0.1657 | 1.125 | 1.457 | .145 |
| Subclassification (Weight supported) | BBB/BMS | 0.6893 | 0.7525 | -0.7855 | 2.164 | 0.9161 | .360 |
| | Paw rotation | 0.9070 | 0.4696 | -0.01348 | 1.827 | 1.931 | .053 |
| Subclassification (Weight supported bipedal treadmill) | BBB/BMS | 0.5781 | 0.1859 | 0.2137 | 0.9425 | 3.109 | .002 |
| | Hindpaw ladder/grid | 0.9908 | 0.6055 | -0.1959 | 2.177 | 1.636 | .102 |
| | Coordination | 0.8355 | 0.6231 | -0.3858 | 2.057 | 1.341 | .180 |
| | Paw rotation | 0.7687 | 0.3555 | 0.07193 | 1.465 | 2.162 | .031 |
| | Stride length | 0.3503 | 0.6820 | -0.9865 | 1.687 | 0.5136 | .608 |
| | ACOS/Tarlov | 1.367 | 0.5553 | 0.2783 | 2.455 | 2.461 | .014 |
| Subclassification (Weight supported quadrupedal treadmill) | BBB/BMS | 0.9012 | 0.2089 | 0.4918 | 1.311 | 4.314 | < .001 |
| | Hindpaw ladder/grid | 0.7801 | 0.5792 | -0.3552 | 1.915 | 1.347 | .178 |
| | Coordination | 1.199 | 1.182 | -1.118 | 3.516 | 1.014 | .311 |
| | Paw rotation | 0.7221 | 0.5160 | -0.2893 | 1.734 | 1.399 | .162 |
| | Stride length | 0.4136 | 1.375 | -2.282 | 3.109 | 0.3007 | .764 |
| Subclassification (Swimming + exercise ball) | Narrow beam | -0.03899 | 0.5774 | -1.171 | 1.093 | -0.06753 | .946 |
| | Hindpaw ladder/grid | 0.6663 | 0.8741 | -1.047 | 2.380 | 0.7622 | .446 |
| | Coordination | -0.6044 | 1.251 | -3.056 | 1.847 | -0.4832 | .629 |
| | Stride length | -0.7677 | 1.412 | -3.536 | 2.000 | -0.5436 | .587 |
| Subclassification (Reaching and Treadmill) | Coordination | 2.103 | 1.317 | -0.4778 | 4.683 | 1.597 | .110 |
| Subclassification (Targeted enrichment + swimming) | Swim score | 2.412 | 1.233 | -0.004549 | 4.829 | 1.956 | .050 |

**Table S7. Continuous meta-regression of total rehabilitation duration and treatment efficacy.**

*Meta-Analytic Tests*

| | Test | *P* |
|---|---|---|
| Residual heterogeneity | $Q_e(209) = 558.33$ | < .001 |
| Pooled effect | z = 8.52 | < .001 |
| Moderation | $Q_m(1) = 4.51$ | .034 |

*Meta-Analytic Estimates*

| | | | 95% CI | | 95% PI | |
|---|---|---|---|---|---|---|
| | Estimate | Standard Error | Lower | Upper | Lower | Upper |
| Pooled effect | 0.8208 | 0.09630 | 0.6320 | 1.010 | 0.3569 | 1.285 |

*Effect Size Meta-Regression Terms Tests*

| | $Q_m$ | df | *P* |
|---|---|---|---|
| Duration total (w) | 4.510 | 1 | .034 |

*Effect Size Meta-Regression Coefficients*

| | | | 95% CI | | | |
|---|---|---|---|---|---|---|
| | Estimate | Standard Error | Lower | Upper | z | *P* |
| Intercept | 0.6710 | 0.1202 | 0.4354 | 0.9066 | 5.581 | < .001 |
| Duration total (w) | 0.02270 | 0.01069 | 0.001751 | 0.04365 | 2.124 | .034 |

**Table S8. Quadratic meta-regression assessing plateauing of rehabilitation efficacy with increasing duration.**

*Meta-Analytic Tests*

| | Test | *P* |
|---|---|---|
| Residual heterogeneity | $Q_e(208) = 557.70$ | < .001 |
| Pooled effect | z = 8.16 | < .001 |
| Moderation | $Q_m(2) = 5.37$ | .068 |

*Meta-Analytic Estimates*

| | | | 95% CI | | 95% PI | |
|---|---|---|---|---|---|---|
| | Estimate | Standard Error | Lower | Upper | Lower | Upper |
| Pooled effect | 0.8211 | 0.1006 | 0.6240 | 1.018 | 0.3270 | 1.315 |

*Meta-Analytic Estimates*

| | Estimate | Standard Error | 95% CI Lower | 95% CI Upper | 95% PI Lower | 95% PI Upper |
|---|---|---|---|---|---|---|

*Effect Size Meta-Regression Terms Tests*

| | $Q_m$ | df | *P* |
|---|---|---|---|
| Duration total (w) | 2.303 | 1 | .129 |
| Duration in weeks squared | 0.9000 | 1 | .343 |

*Note.* Fixed effects tested using z-distribution.

*Effect Size Meta-Regression Coefficients*

| | Estimate | Standard Error | 95% CI Lower | 95% CI Upper | z | *P* |
|---|---|---|---|---|---|---|
| Intercept | 0.5652 | 0.1696 | 0.2329 | 0.8976 | 3.334 | < .001 |
| Duration total (w) | 0.05637 | 0.03714 | -0.01644 | 0.1292 | 1.517 | .129 |
| Duration in weeks squared | -0.002111 | 0.002225 | -0.006473 | 0.002250 | -0.9487 | .343 |

#### Table S9. Intercept and no-intercept meta-regression models of rehabilitation onset phase.

*Meta-Analytic Tests*

| | Test | *P* |
|---|---|---|
| Residual heterogeneity | $Q_e(204) = 550.17$ | < .001 |
| Pooled effect | z = 4.18 | < .001 |
| Moderation | $Q_m(4) = 4.34$ | .362 |

*Meta-Analytic Estimates*

| | Estimate | Standard Error | 95% CI Lower | 95% CI Upper | 95% PI Lower | 95% PI Upper |
|---|---|---|---|---|---|---|
| Pooled effect | 0.6468 | 0.1546 | 0.3438 | 0.9499 | -0.3117 | 1.605 |

*Effect Size Meta-Regression Terms Tests*

| | $Q_m$ | df | *P* |
|---|---|---|---|
| Onset phase | 4.336 | 4 | .362 |

*Effect Size Meta-Regression Coefficients*

| | Estimate | Standard Error | 95% CI Lower | 95% CI Upper | z | *P* |
|---|---|---|---|---|---|---|
| Intercept | 0.5642 | 0.1626 | 0.2454 | 0.8829 | 3.469 | < .001 |
| Onset phase (2_early subacute) | 0.07589 | 0.09408 | -0.1085 | 0.2603 | 0.8066 | .420 |

*Effect Size Meta-Regression Coefficients*

| | Estimate | Standard Error | 95% CI Lower | 95% CI Upper | z | *P* |
|---|---|---|---|---|---|---|
| Onset phase (3_late subacute) | 0.1935 | 0.1131 | -0.02812 | 0.4152 | 1.711 | .087 |
| Onset phase (4_intermediate) | 0.2392 | 0.1539 | -0.06236 | 0.5409 | 1.555 | .120 |
| Onset phase (5_chronic) | 0.06999 | 0.2066 | -0.3349 | 0.4748 | 0.3388 | .735 |

*Effect Size Meta-Regression Coefficients* (no intercept)

| | Estimate | Standard Error | 95% CI Lower | 95% CI Upper | z | *P* |
|---|---|---|---|---|---|---|
| Onset phase (1_acute) | 0.5642 | 0.1626 | 0.2454 | 0.8829 | 3.469 | < .001 |
| Onset phase (2_early subacute) | 0.6400 | 0.1627 | 0.3211 | 0.9590 | 3.933 | < .001 |
| Onset phase (3_late subacute) | 0.7577 | 0.1752 | 0.4143 | 1.101 | 4.325 | < .001 |
| Onset phase (4_intermediate) | 0.8034 | 0.2069 | 0.3979 | 1.209 | 3.883 | < .001 |
| Onset phase (5_chronic) | 0.6341 | 0.2478 | 0.1484 | 1.120 | 2.559 | .011 |

#### Table S10. Meta-regression of daily rehabilitation exposure and treatment efficacy.

*Meta-Analytic Tests*

| | Test | *P* |
|---|---|---|
| Residual heterogeneity | $Q_e(190) = 514.17$ | < .001 |
| Pooled effect | z = 7.21 | < .001 |
| Moderation | $Q_m(4) = 15.07$ | .005 |

*Meta-Analytic Estimates*

| | Estimate | Standard Error | 95% CI Lower | 95% CI Upper | 95% PI Lower | 95% PI Upper |
|---|---|---|---|---|---|---|
| Pooled effect | 0.8272 | 0.1147 | 0.6023 | 1.052 | 0.2324 | 1.422 |

*Effect Size Meta-Regression Terms Tests*

| | $Q_m$ | df | *P* |
|---|---|---|---|
| Timeframe spent in rehab | 15.07 | 4 | .005 |

*Effect Size Meta-Regression Coefficients*

| | Estimate | Standard Error | 95% CI Lower | 95% CI Upper | z | *P* |
|---|---|---|---|---|---|---|
| Intercept (≤10 min) | 0.8950 | 0.1490 | 0.6030 | 1.187 | 6.007 | < .001 |
| 11-30 min | -0.1565 | 0.1157 | -0.3833 | 0.07032 | -1.352 | .176 |
| 31-59 min | -0.3387 | 0.1501 | -0.6329 | -0.04463 | -2.257 | .024 |

*Effect Size Meta-Regression Coefficients*

| | Estimate | Standard Error | 95% CI Lower | 95% CI Upper | z | *P* |
|---|---|---|---|---|---|---|
| ≥60 min | 0.06492 | 0.1508 | -0.2306 | 0.3604 | 0.4306 | .667 |
| *Ad libitum* | 0.1119 | 0.1318 | -0.1464 | 0.3703 | 0.8492 | .396 |

*Effect Size Meta-Regression Terms Tests*

| | $Q_m$ | df | *P* |
|---|---|---|---|
| Timeframe spent in rehab | 66.39 | 5 | < .001 |

*Effect Size Meta-Regression Coefficients* (no intercept)

| | Estimate | Standard Error | 95% CI Lower | 95% CI Upper | z | p |
|---|---|---|---|---|---|---|
| ≤10 min | 0.8950 | 0.1490 | 0.6030 | 1.187 | 6.007 | < .001 |
| 11-30 min | 0.7385 | 0.1211 | 0.5010 | 0.9759 | 6.096 | < .001 |
| 31-59 min | 0.5562 | 0.1569 | 0.2487 | 0.8638 | 3.545 | < .001 |
| ≥60 min | 0.9599 | 0.1579 | 0.6504 | 1.269 | 6.080 | < .001 |
| *Ad libitum* | 1.007 | 0.1358 | 0.7407 | 1.273 | 7.414 | < .001 |

#### Table S11. Daily rehabilitation exposure differentially influences forced and voluntary rehabilitation efficacy.

*Effect Size Meta-Regression Terms Tests*

| | Subgroup | $Q_m$ | df | *P* |
|---|---|---|---|---|
| Timeframe spent in rehab | Forced | 6.526 | 3 | .089 |
| | Voluntary | 5.691 | 2 | .058 |

.

*Effect Size Meta-Regression Coefficients*

| | Subgroup | Estimate | Standard Error | 95% CI Lower | 95% CI Upper | z | *P* |
|---|---|---|---|---|---|---|---|
| Intercept (≤10 min) | Forced | 0.9399 | 0.1079 | 0.7284 | 1.151 | 8.712 | < .001 |
| | Voluntary | 0.5541 | 0.3093 | -0.05219 | 1.160 | 1.791 | .073 |
| 11-30 min | Forced | -0.2423 | 0.1214 | -0.4802 | -0.004384 | -1.996 | .046 |
| 31-59 min | Forced | -0.3804 | 0.1535 | -0.6812 | -0.07948 | -2.478 | .013 |
| ≥60 min | Forced | -0.2606 | 0.1842 | -0.6215 | 0.1004 | -1.415 | .157 |
| | Voluntary | 0.7280 | 0.3087 | 0.1229 | 1.333 | 2.358 | .018 |
| *Ad libitum* | Voluntary | 0.4549 | 0.2770 | -0.08812 | 0.9979 | 1.642 | .101 |

#### Table S12. Continuous meta-regression of daily forced rehabilitation duration and treatment efficacy.

*Meta-Analytic Tests*

| | Test | *P* |
|---|---|---|
| Residual heterogeneity | $Q_e(140) = 347.30$ | < .001 |
| Pooled effect | z = 16.60 | < .001 |
| Moderation | $Q_m(1) = 6.57$ | .010 |

*Meta-Analytic Estimates*

| | | | 95% CI | | 95% PI | |
|---|---|---|---|---|---|---|
| | Estimate | Standard Error | Lower | Upper | Lower | Upper |
| Pooled effect | 0.7042 | 0.04243 | 0.6210 | 0.7873 | 0.6210 | 0.7873 |

*Effect Size Meta-Regression Terms Tests*

| | $Q_m$ | df | *P* |
|---|---|---|---|
| Duration per day (m) | 6.567 | 1 | .010 |

*Effect Size Meta-Regression Coefficients*

| | | | 95% CI | | | |
|---|---|---|---|---|---|---|
| | Estimate | Standard Error | Lower | Upper | z | *P* |
| Intercept | 0.9355 | 0.09741 | 0.7446 | 1.126 | 9.604 | < .001 |
| Duration per day (m) | -0.007774 | 0.003034 | -0.01372 | -0.001828 | -2.563 | .010 |

### Supplementary Figures

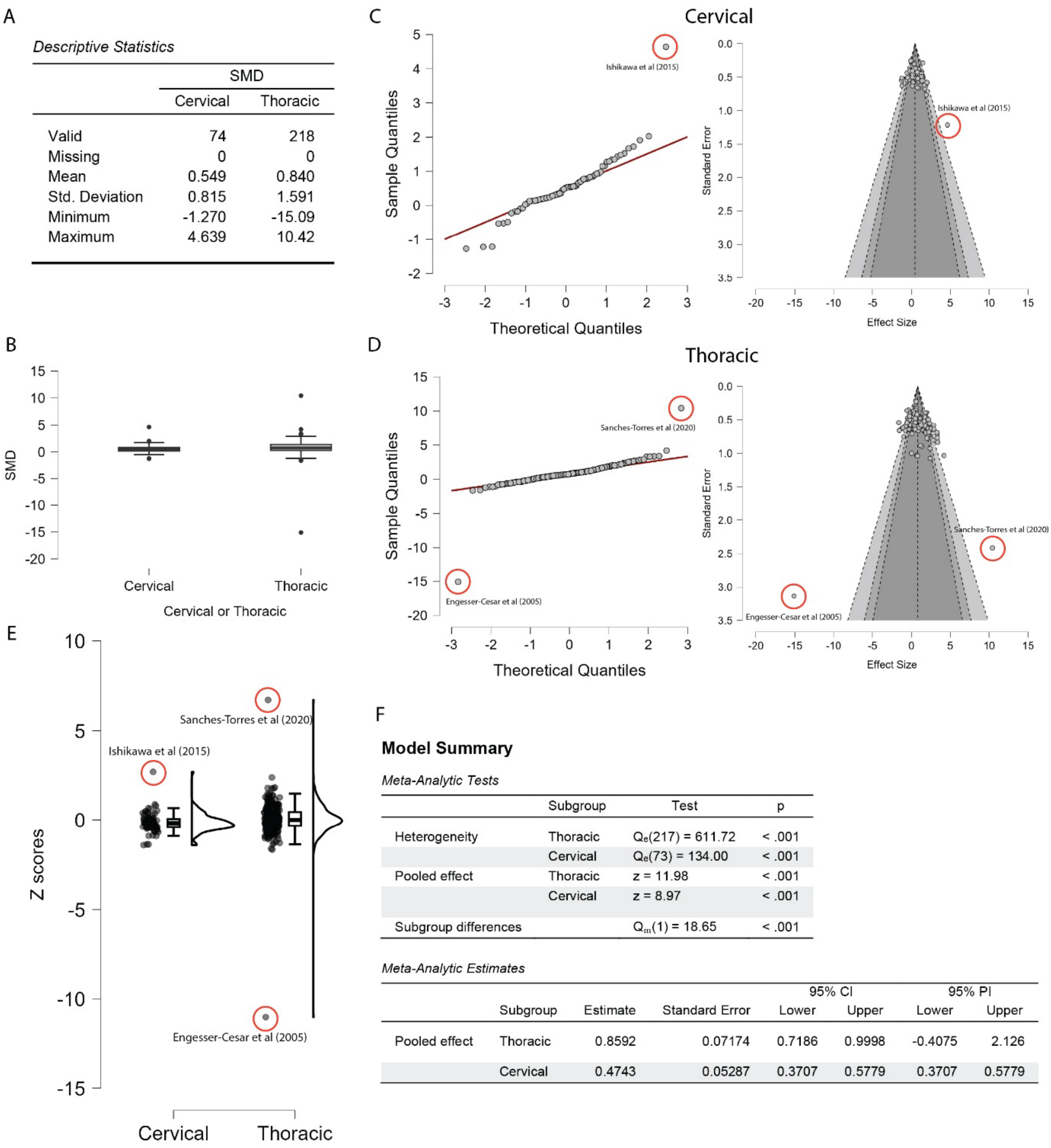


*Descriptive Statistics*

| | SMD | |
|---|---|---|
| | Cervical | Thoracic |
| Valid | 74 | 218 |
| Missing | 0 | 0 |
| Mean | 0.549 | 0.840 |
| Std. Deviation | 0.815 | 1.591 |
| Minimum | -1.270 | -15.09 |
| Maximum | 4.639 | 10.42 |

**Model Summary**

*Meta-Analytic Tests*

| | Subgroup | Test | p |
|---|---|---|---|
| Heterogeneity | Thoracic | $Q_e(217) = 611.72$ | < .001 |
| | Cervical | $Q_e(73) = 134.00$ | < .001 |
| Pooled effect | Thoracic | z = 11.98 | < .001 |
| | Cervical | z = 8.97 | < .001 |
| Subgroup differences | | $Q_m(1) = 18.65$ | < .001 |

*Meta-Analytic Estimates*

| | | | | 95% CI | | 95% PI | |
|---|---|---|---|---|---|---|---|
| | Subgroup | Estimate | Standard Error | Lower | Upper | Lower | Upper |
| Pooled effect | Thoracic | 0.8592 | 0.07174 | 0.7186 | 0.9998 | -0.4075 | 2.126 |
| | Cervical | 0.4743 | 0.05287 | 0.3707 | 0.5779 | 0.3707 | 0.5779 |

**Fig. S1. Outlier detection and distribution analysis across cervical and thoracic injury datasets. (A)** Summary statistics for cervical and thoracic injury datasets, including the number of studies, mean, standard deviation, minimum, and maximum values. **(B)** Boxplots illustrating the distribution and variability within each dataset. **(C)** Normal Q-Q plot and funnel plot for the cervical dataset, identifying a single outlier. **(D)** Normal Q-Q plot and funnel plot for the thoracic dataset, indicating multiple outliers. **(E)** Raincloud plots of Z-scores calculated for each study; scores exceeding 2.5 were classified as outliers. Three studies are highlighted. **(F)** Model summary statistics from the REML meta-analysis, stratified by injury level.

A

*Descriptive Statistics*

| | SMD | |
|---|---|---|
| | Cervical | Thoracic |
| Valid | 73 | 216 |
| Missing | 0 | 0 |
| Mean | 0.493 | 0.869 |
| Std. Deviation | 0.662 | 0.972 |
| Minimum | -1.270 | -1.625 |
| Maximum | 2.028 | 4.190 |

B

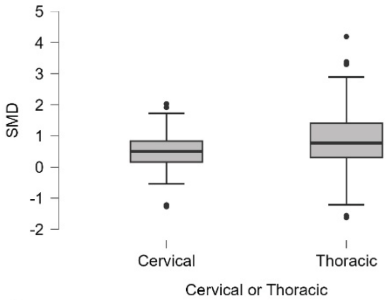


C

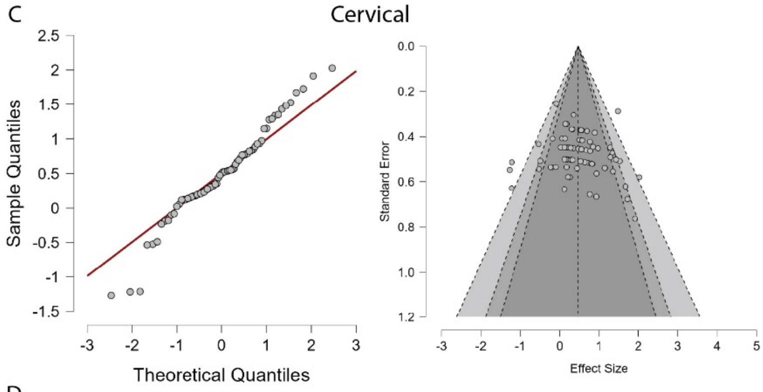


D

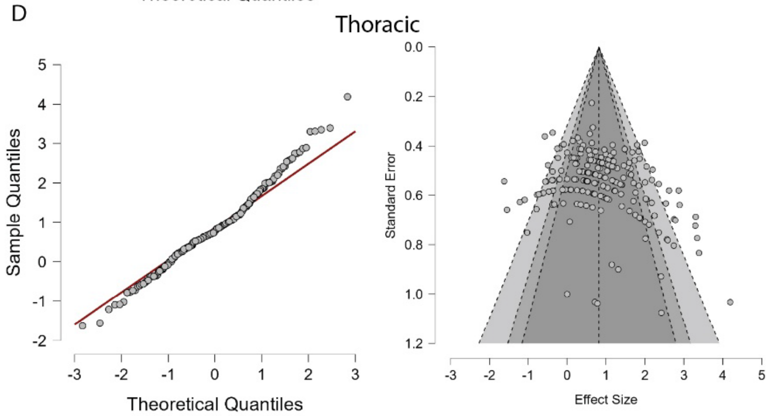


E

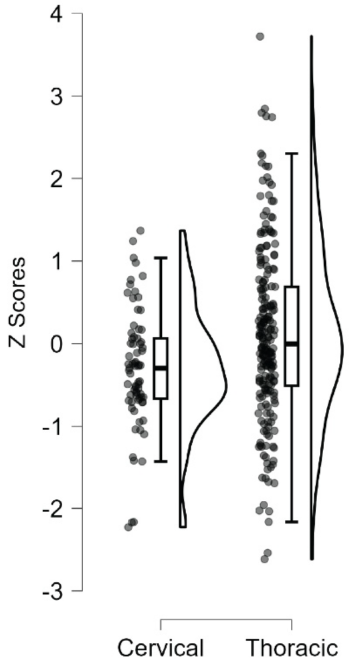


F

**Model Summary**

*Meta-Analytic Tests*

| | Subgroup | Test | p |
|---|---|---|---|
| Heterogeneity | Thoracic | $Q_e(215) = 570.13$ | < .001 |
| | Cervical | $Q_e(72) = 122.22$ | < .001 |
| Pooled effect | Thoracic | z = 12.05 | < .001 |
| | Cervical | z = 8.81 | < .001 |
| Subgroup differences | | $Q_m(1) = 19.17$ | < .001 |

*Meta-Analytic Estimates*

| | | | | 95% CI | | 95% PI | |
|---|---|---|---|---|---|---|---|
| | Subgroup | Estimate | Standard Error | Lower | Upper | Lower | Upper |
| Pooled effect | Thoracic | 0.8536 | 0.07087 | 0.7147 | 0.9925 | -0.3957 | 2.103 |
| | Cervical | 0.4664 | 0.05292 | 0.3627 | 0.5701 | 0.3627 | 0.5701 |

**Fig. S2. Descriptive statistics and distribution analysis of the cervical and thoracic injury datasets following outlier removal. (A)** Summary statistics for cervical and thoracic injury datasets, including the number of studies, mean, standard deviation, minimum, and maximum values. **(B)** Boxplots illustrating the distribution and variability within each dataset. **(C)** Normal Q-Q plot and funnel plot for the cervical dataset. **(D)** Normal Q-Q plot and funnel plot for the thoracic dataset. **(E)** Raincloud plots of Z-scores calculated for each study; scores exceeding 2.5 were classified as outliers. **(F)** Model summary statistics from the REML meta-analysis, stratified by injury level.

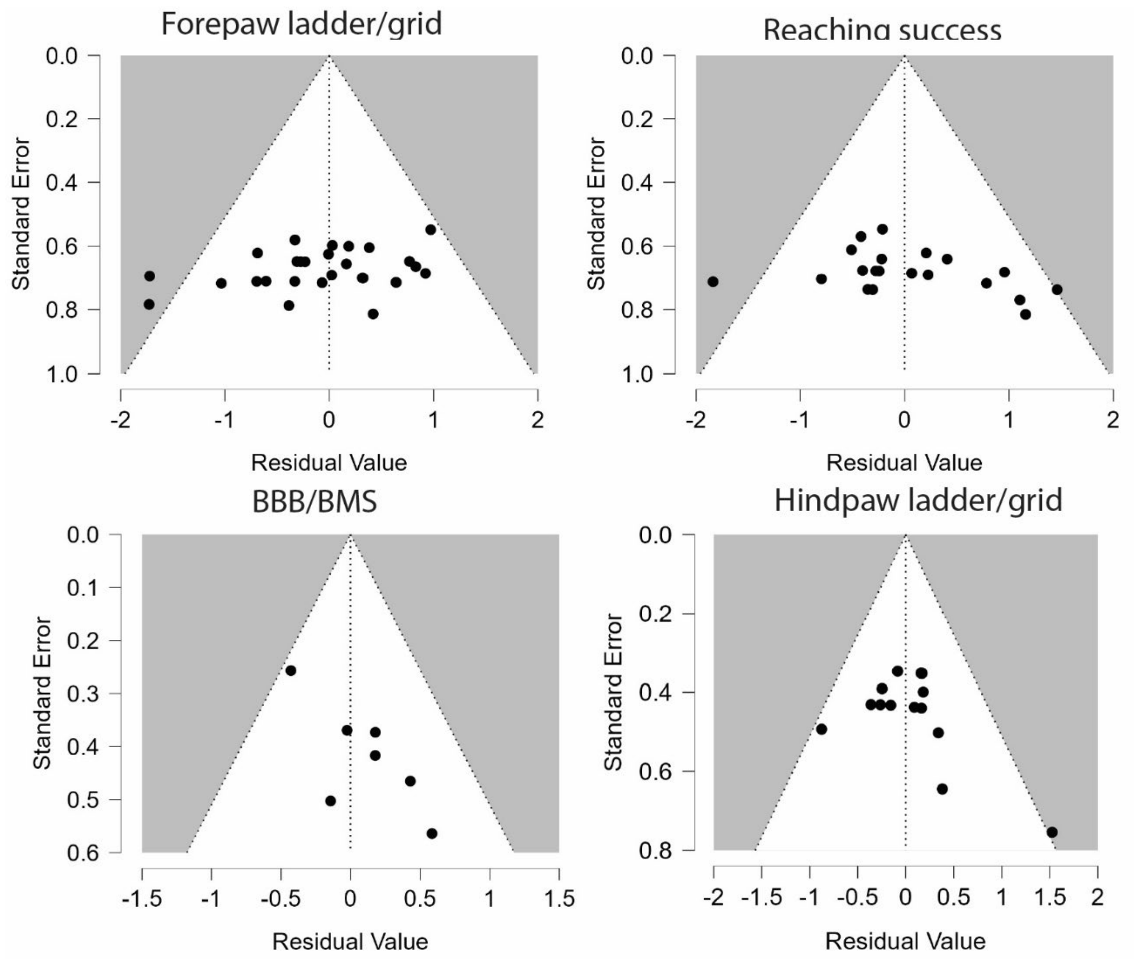


**Fig. S3. Residual funnel plots for the cervical dataset stratified by behavior measure.** Residuals (observed minus predicted effect sizes) are plotted on the horizontal axis against study precision (Standard Error) on the vertical axis. The dashed vertical line represents a residual of zero. The slanted dashed lines indicate the expected 95% pseudo-confidence intervals. The asymmetrical distribution of studies in the lower-left quadrant suggests potential publication bias or the omission of a significant study-level covariate.

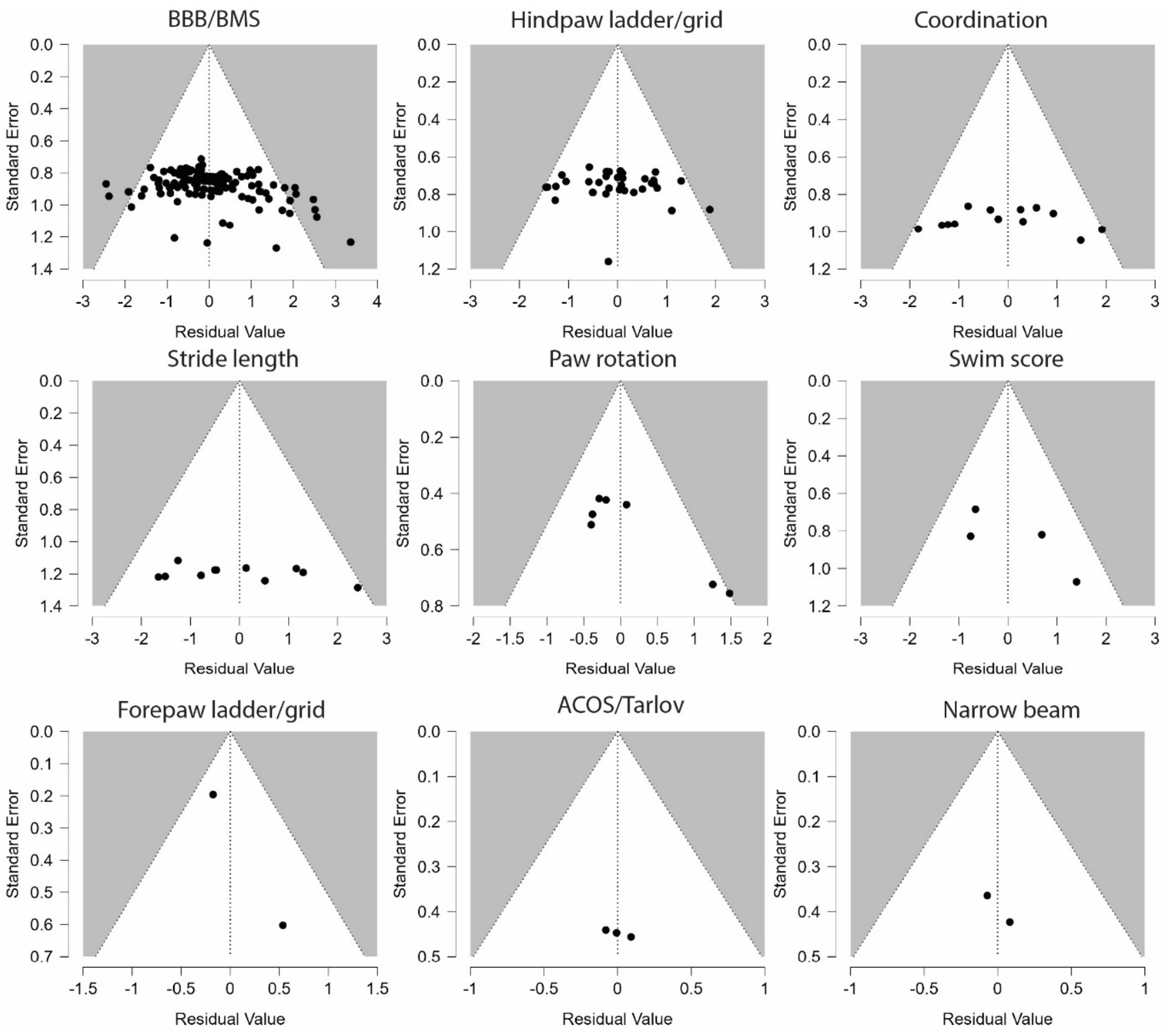


**Fig. S4. Residual funnel plots for the thoracic dataset stratified by behavior measure.** Residuals (observed minus predicted effect sizes) are plotted on the horizontal axis against study precision (Standard Error) on the vertical axis. The dashed vertical line represents a residual of zero. The slanted dashed lines indicate the expected 95% pseudo-confidence intervals. The asymmetrical distribution of studies in the lower-left quadrant suggests potential publication bias or the omission of a significant study-level covariate.

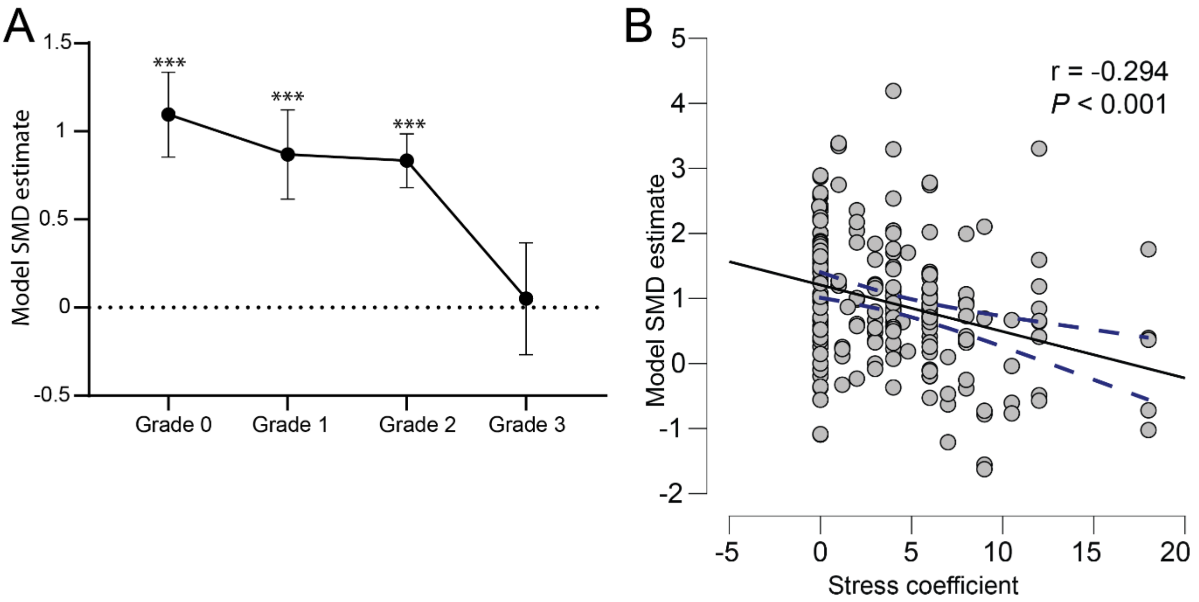


**Fig. S5. The association between animal stress on the model estimates and the effect of rehabilitation. (A)** A stress grade was assigned to each rehabilitation type based on its estimated propensity to elicit physiological **st**ress in the animals. (B) This grade was multiplied by the duration spent in the rehabilitation, resulting in a 'stress coefficient'. Model-derived SMD estimates were then correlated with the stress coefficient using Pearson's correlati*on*. The correlation coefficient and P value are reported.
